# Dpp and Hedgehog promote the Glial response to neuronal damage in the developing Drosophila Visual system

**DOI:** 10.1101/2020.02.24.960625

**Authors:** Sergio B. Velarde, Alvaro Quevedo, Carlos Estella, Antonio Baonza

## Abstract

Damage in the nervous system induces a stereotypical response that is mediated by glial cells. Here, we use the eye disc to explore the mechanisms involved in promoting glial cell response after neural injuries. We demonstrate that eye glia cells rapidly respond to neuronal injury by increasing in number and undergoing morphological changes, which grant them phagocytic abilities. We found that this glial response is controlled by the activity of the long-range signalling pathways, *decapentaplegic* (dpp) and *hedgehog* (hh). These pathways are activated in the damaged region and their functions are necessary for inducing glial cell proliferation and migration to the eye discs. The latter of these two processes depends on the function of the JNK pathway, which is cooperatively activated by *dpp* and *hh* signalling.

## INTRODUCTION

A complex nervous system comprises of neurons and glial cells whose development and function are mutually interdependent. The intricate interaction between these two cell types is essential for the generation and maintenance of a functional nervous system. During development or after neural injuries, cells within the nervous system accommodate changes in order to preserve structural integrity and function. Glial cells actively participate in all aspects of nervous system development, including the mechanism involved in maintaining structural robustness and functional plasticity. In response to neuronal damage, glial cells proliferate, change their morphology and alter their behaviour (Kato et al., 2018; Klambt, 2009; Brosius Lutz and Barres, 2014). This glial cell response is associated with their regenerative function and is found across species. The signalling pathways underlying glial response and how they are coordinated; however, remains poorly understood, but progress can be made to build our understanding of these cellular processes by exploring how glial cells respond to injury.

*Drosophila melanogaster* is an excellent model system to discover evolutionarily conserved gene functions and gene networks. Previously, the eye disc of *Drosophila* has been used as a model system to analyse the basic mechanisms regulating the migration of glial cells along their neuronal partners (Rangarajan et al., 1999; Franzdottir et al., 2009; Yildirim et al., 2019; Yuva-Aydemir and Klambt, 2011; Chotard and Salecker, 2007; Silies et al., 2007; Silies et al., 2010; Klambt, 2009). The eye disc develops from a group of ectodermal cells with embryonic origin, distinct from the neuroectodermal cells that form the central nervous system (CNS). Thus, unlike its mammalian counterpart, *Drosophila* eyes are not part of the CNS. Nevertheless, similarly to mammalian systems, they do contain neurons and glial cells. The eye primordia develops progressively, from posterior to anterior, over the course of approximately two days. A furrow (known as morphogenetic furrow (MF)) sweeps across the disc during this period, leaving in its wake developing clusters of photoreceptor cells that will become the individual units of the compound eye, known as ommatidia (Wolff and Ready, 1993). Therefore, the region anterior to the furrow is mainly constituted of proliferating, undifferentiated cells, whereas the region posterior to the MF consists predominantly of cells which have exited the cell cycle and have begun to differentiate into photoreceptors (Wolff and Ready, 1993; Baker and Yu, 2001; Baonza and Freeman, 2005). Unlike photoreceptors, the progenitors of all subretinal glia are not generated from the eye disc cells. The developing imaginal disc is connected to the brain by the optic stalk. During early embryonic stages, the anlage of the eye disc is established and a few glial cells are born in the initial segment of the Bolwing nerve, which will later become the optic stalk (Chotard and Salecker, 2007; Choi and Benzer, 1994; Silies et al., 2007). These glial cells are the precursors of the eye disc glial cells. During larval stages, these precursor cells proliferate forming new glial cells that accumulate in the optic stalk. As the eye imaginal disc grows and neurogenesis is initiated behind the MF, glial cells leave the optic stalk and migrate onto the eye disc (Chotard and Salecker, 200; Choi and Benzer, 1994; Silies et al., 2007; Silies et al., 2010).

The eye discs contain distinct glial cell types (Silies et al., 2007). Subperineurial cells, the so-called carpet cells, are two large cells that cover the entire differentiated part of the eye disc epithelium. Sitting basally to these two cells are the perineurial glial cells that have a distinct morphology in the optic stalk and in the eye disc (Silies et al., 2007; Sasse and Klambt, 2016). These cells define a reserve pool, which can generate glial cells when necessary (for plasticity, and development); accordingly, these cells maintain the ability to divide during eye disc development (Silies et al., 2007). Carpet cells separate the perineurial glia from the underlying wrapping glia. Wrapping glial cells derive from perineurial glial and enwrap all axons produced by the photoreceptors. The wrapping glial cells perform functions which resemble the non-myelinating Schwann cells forming Remak fibers in the mammalian peripheral nervous system (PNS) (Yildirim et al., 2019). Schwann cells play a key role in promoting regeneration and provide the high ability of the peripheral nerves to regenerate (Jessen and Mirsky, 2016).

Many studies using eye discs as a model system have contributed to uncovering the basic mechanisms and signalling pathways involved in the coupling of neuronal and glial development, and have shown that *Drosophila* glia can serve as an experimental model to gain insights into mammalian glial biology (Franzdottir et al., 2009; Sasse et al., 2015; Silies et al., 2007; Yildirim et al., 2019). However, despite the widespread use of this model to study the mechanisms that regulate the coordinated development of neurons and glial cells, little is known about the response of glial cells to damage and the signalling pathways that might be mediating this function. The unique developmental features of the fly eye disc, make this structure an excellent model to analyse the signals emitted by damaged neurons and the mechanism involved in regulating glial cell response. In addition, considering the similitudes between non**-**myelinating Schwann cells and wrapping glial, the eye discs might provide new insights into the signalling pathway network involved in regulate glial cell behaviour in response to damage in the PNS.

Here, we use the eye disc to explore the mechanisms involved in promoting glial cell response to neural injuries during the development. We demonstrate that eye glia cells rapidly respond to neuronal injury by increasing in number and underging morphological changes that confer to them phagocytic activity. We found that this glial response is controlled by the activity of the long-range signalling pathways, *decapentaplegic* (dpp) and *hedgehog* (hh). These pathways are activated in the damaged region and their function is necessary for inducing the proliferation and migration of glial cells to the eye discs. This latter process depends on the function of the JNK pathway, which is cooperatively activated by *dpp* and *hh* signalling.

## RESULTS

### Cell death induction in the eye disc epithelium promotes the accumulation of glial cells

Different studies have shown that when the retinal region of eye discs is damaged, a regenerative response is initiated that includes compensatory proliferation (Fan and Bergmann, 2008). However, whether this response also involves glia cell activation, as it occurs in other nervous system region, was unknown. In order to study glial cell response to damage in the eye disc, we have examined the localization, pattern of proliferation and number of glia cells in eye discs that have been damaged in the retinal region. To genetically induce targeted damage in this region of the eye disc epithelium, we used the *Gal4/UAS/Gal80^Ts^* system to transiently over-express the pro-apoptotic gene *reaper* (*rpr*) under the control of the eye specific *glass multiple reporter* (*GMR*) *Gal4* line. *GMR-Gal4* is expressed in the eye disc, in all cells posterior to the morphogenetic furrow (MF), including photoreceptor cells. Given that, as we mentioned above, glial cells are not originated in the eye disc, this driver is not active in glial cells (Figure 1). Therefore, any change in the behaviour of glial cells would be non-autonomously induced by apoptotic cells in the retinal region. We used the *tub-Gal80^ts^* transgene to modulate the time of apoptotic induction, thus *GMR-Gal4 tub-Gal80^ts^ UAS-rpr* larvae were raised at the permissive temperature and then shifting to restrictive temperature for three days to over-express *UAS-rpr* (see methods). As expected, we found that cell death strongly increases behind the MF in these discs, as assayed by staining for the apoptotic marker, anti-cleaved Dcp1 (Figure S1). Remarkably, we observed a pronounced increase in the density of glial cells in damaged discs compared to control discs (0.018±0.0004 vs 0.009±0.0003 in control discs) (Figure 1 D and G and supplementary Table). Furthermore, in contrast to control discs, where glial cells are always located on the basal layer of the discs (Figure 1 C-C’’’’), in injured discs we found glial cells contacting with photoreceptors in the middle layer of the eye disc epithelium (yellow arrows in Figure 1 D’’-D’’’’). We also observed a number of glia cells located apically to the photoreceptors (green arrows in Figure 1 D-D’’’’). Interestingly, retinal damage not only affects the apical/basal location of glia, but also their positioning with respect to the MF. In control discs, the anterior border of glial migration lies between 0**–**5 cell rows behind the most anterior row of photoreceptors (Figure 1 E-E’ and H). However, in damaged discs, the anterior boundary of the glial migration moved ahead, with some glia even migrating beyond the morphogenetic furrow (Figure 1 F, F’ and H).

**Figure 1.**
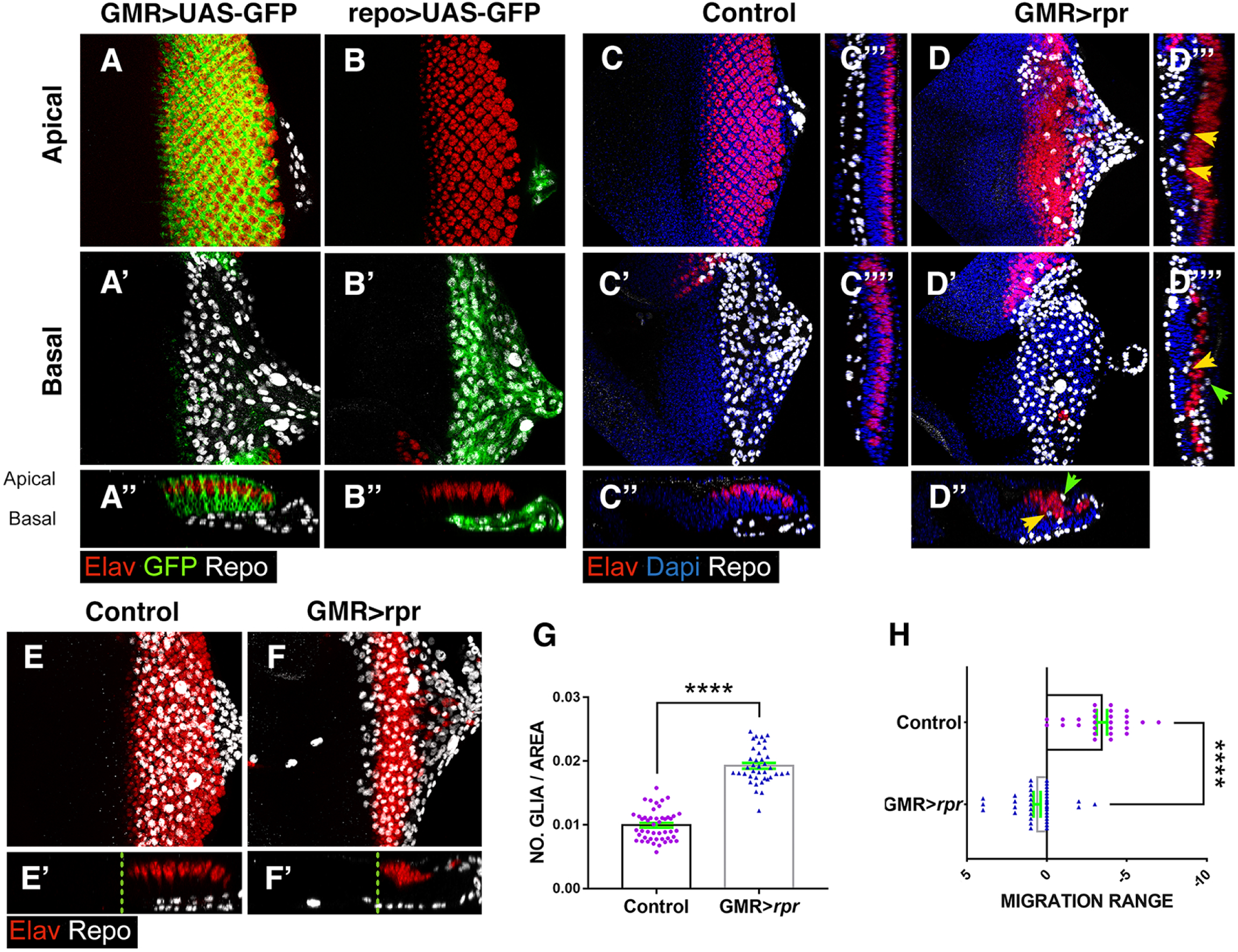
Response of subretinal glia to neuronal damage in the eye discs. (A-F) Third instar eye discs stained with anti-Elav (Photorreceptors in red), anti-Repo (glial cells in white) and the nuclear marker Dapi (blue). (A-D) Apical/middle layers of the eye disc ephilelium. (A’-D’) Basal layer of the eye discs. (A’’-F’) The X–Z projections show cross-sections of the eye discs shown in A-F perpendicular to the furrow. (C’’’-C’’’’ and D’’’-D’’’’) Transverse sections perpendicular to the furrow of discs shown in C-D. (A-A’’) Eye disc showing the expression of *UAS-GFP* (green) under the control of *GMR-Gal4*. Glia nuclei (anti-Repo in white) are located in the basal layer of the eye disc and they do not express *UAS-GFP*. (B-B’’) The expression of *UAS-GFP* (green) under the control of *repo–Gal4* is restricted to glia cells (B’-B’’). (C-C’’’’) In control discs subretinal glia cells are always located in the basal layer of the disc. (D-D’’’’) In damaged *UAS-rpr/+ GMR-Gal4, tub-Gal80^ts^* eye discs glial cells are not only in the basal layer of the disc but also in the middle and apical layers. This is most clearly seen in transverse sections shown in (D’’-D’’’’). Arrowheads indicate glia cells located in apical (green) and middle (yellow) layers of the discs (D’’-D’’’’). Note that glia in medial layer directly contact with photorreceptors. (E-F’) Third instar control (*GMR-Gal4 tub-Gal80^ts^*) (E) and damaged (*UAS-rpr/+ GMR-Gal4, tub-Gal80^ts^*) (F) eye showing the relative position of the anterior border of glial migration (dotted line) with respect to the anteriormost row of photoreceptors. In damaged discs the anterior border of glial migration lies 2–4 row in front of the morphogenetic furrow. (G) Graph shows the density of glia cells (Number of glia cells/area occupy by glia) in control and damaged *UAS-rpr/+ GMR-Gal4 tub-Gal80^ts^* discs. (H) Graph shows the relative position of the anterior border of glial migration with respect to the anteriormost row of photoreceptors (0 indicates the position of this row) in control and damaged *UAS-rpr/+ GMR-Gal4 tub-Gal80^ts^* eye discs. Here, and in the rest of the figures statistical analysis is shown in supplementary Table. The error bars represent SEM.

In order to get a better understanding of the sequence of events associated with subretinal glia response, we examined glia behaviour at different times after retinal damage. To that end, *GMR-Gal4 tub-Gal80^ts^ UAS-rpr* larvae were raised at permissive temperature and then shifted to the restrictive temp for 24 hours to induce *UAS-rpr* (T0) and then shifted back to the permissive temperature to allow time for them to recover (see method). Discs were then analysed at different timepoints (Figure 2). We found that the ectopic expression of *UAS*-*rpr* for 24 hours resulted in massive apoptosis of cells located behind the MF (Figure 2 C-D’’). Most apoptotic cells were located at the basal region of the discs, and predominantly corresponded to interommatidial cells, as they did not express the neuronal marker Elav (Figure 2 C-D’’). In discs dissected immediately after genetic ablation (T0), we observed that the density of glia cells was already significantly higher than in control discs (Figure 2 C-D’’ I and J). This is mainly due to the increased number of glial cells that appear in the middle/apical layers of the disc epithelium (Figure 2 J). After 24 hours (T1) of recovery at the permissive temperature, most cellular debris was displaced toward the posterior region of the discs and even inside the optic stalk (Figure 2 E-F’’). At this time the density of glia cells was still higher in damaged discs compared to control discs; however, numbers were similar to those found at T0 (Figure 2 I and K). We also observed a high number of glia cells in the middle/apical layers of the retinal epithelium (Figure 2 E-F’’). After 48hs of recovery (T2), we did not see apoptotic cells or cellular debris in the epithelium. The density of glial cells was similar to the earlier time points, but higher when compared to control discs (Figure 2G-H’’, I and L). However, in contrast to the discs analysed at T0 and T1, we did not find glial cells in the middle or apical layers. In the posterior region of the disc we observed a band of disorganized photoreceptors localized in the basal layer of the discs, that likely correspond to the photoreceptors that did no die after over-expressing *UAS-rpr* (Figure 2G-H’’). The adult eyes derived from these discs were only slightly smaller than control eyes, and they did not show any detectable pattern defect, we occasionally observed small scars in the posterior region of the eye (Figure 2 M-O).

**Figure 2.**
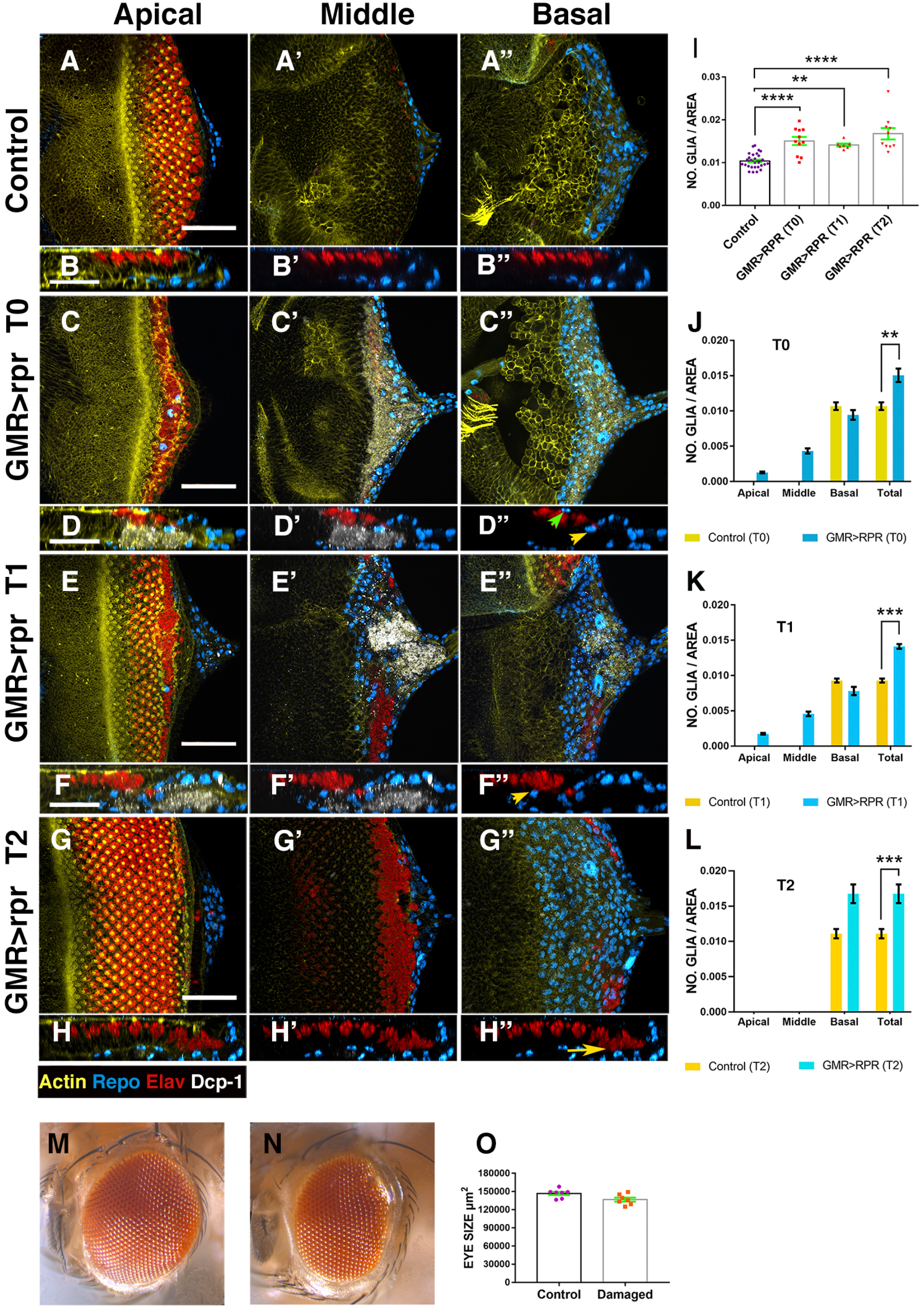
Overview of Glial response at different times after damaging eye. (A-H’’) Third instar eye discs stained with anti-Elav (red) and anti-Repo (blue), the apoptotic marker Dcp-1 (grey) and Phalloidin to visualize F-actina (yellow). Control undamaged disc (*GMR-Gal4 tub-Gal80^ts^*) (A-B’’) and *UAS-rpr/+; GMR-Gal4 tub-Gal80^ts^* eye discs analysed at different times after damaging (C-H’’). (A, C, E and G) Apical, (A’, C’, E’ and G’) middle and (A’’, C’’, E’’ and G’’) basal layers of the eye disc ephilelium. (B-B’’, D-D’’, F-F’’ and H-H’’) X–Z projections show a cross-section of the eye discs epithelium perpendicular to the furrow of the discs shown in A (B-B’’), C (D-D’’), E (F-F’’) and G (H-H’’). (I) Bar charts show the average density of glial cells of discs analysed at different times after damaged induction. (J-L) Bar charts show the average density of glial cells and their apical/basal localization of discs immediately after apoptotic induction T0 (J), discs analysed after 24 h of recovering T1 (K) and discs examined after 48 h of recovering T2 (L). For each timepoints are shown the density of glia in the Apical, middle, and basal layers of the discs, as well as the total density of glial cells for control (*GMR-Gal4 tub-Gal80^ts^*) and *UAS-rpr/+; GMR-Gal4 tub-Gal80^ts^* eye discs. (A-B’’’) In control discs subretinal glial cells, are always localized in the basal layer of the eye disc. (C-D’’) *UAS-rpr/+; GMR-Gal4 tub-Gal80^ts^* eye discs dissected and analysed immediately after ablation (T0). Apoptotic cells are located in the middle and basal layer of the discs, whereas glial cells appear in the middle (yellow arrowhead) as well as apical planes (green arrowhead) (D-D’’). (E-F’’) *UAS-rpr/+; GMR-Gal4 tub-Gal80^ts^* eye discs analysed after 24 h of recovery. The new rows of photoreceptors that are specified during the recovery time are not affected. The average density of glial cells in these discs is still higher than in control discs (K). We find a high number of glial cells in apical and middle layers of the discs (yellow arrowhead, F-F’’). (G-H’’) Damaged eye discs analysed after 48 h of recovery. In these discs we do not find apoptotic debris (in white) (H-H’’). In the posterior region of the disc we observed very disorganized photoreceptors, which are located in the basal layer (yellow arrow in F’’). (M-N) Adult control eye (M) and an eye derived from damaged eye discs (N). (O) Bar charts show the average size of control eyes and adult eye develop from damaged discs. Statistical analysis is shown in supplementary Table. Scale bar 50μm

All together our data suggest that retinal damage activates a non-autonomous response in subretinal glial cells located in the basal layer of the eye disc epithelium. Moreover, our data indicates that developing eye discs have the ability to recover after being damaged for a limited period of time.

### Glia cell proliferation increases in response to retinal damage

The increased number of glial cells observed in damaged eye discs may be due to overmigration of these cells from the optic stalk and/or an excess of glial proliferation. To distinguish between these alternatives, we next examined the proliferation pattern of the subretinal glia in *GMR-Gal4 tub-Gal80^ts^ UAS-rpr* eye discs damaged over a 72 hour period. We observed that in these damaged eye discs, the proportion of glial cells in S phase was higher than in age-matched control animals, as assayed by 5-Ethynyl-20-deoxyuridine (EdU) incorporation (Figure S2 A-B’’ and E). In addition, we also observed an overall increase in the number of dividing glia upon damage, thus the percentage of dividing glia cells increases in damaged eye discs versus control conditions (Figure S2). Only glia cells located in the basal layer of eye disc undergoing mitosis, as we do not find cells expressing PH3 outside this level (Figure S2 C-D’’).

Altogether, our results suggest that in response to damage, the number of glial cells that progress in the cycle and proliferate increases when compared to undamaging conditions.

Following this, we analysed the proliferation dynamics of subretinal glia cells during the recovery time. To this end, eye discs were damaged during a 24 hour period and the number of positive PH3 glial cells was determined at various timepoints post damage induction. We found that at T0 (immediately after damage induction) glia cell proliferation was already elevated compared to control discs. After 24 hours of recovery (T1) glia proliferation was still higher than in undamaged discs, but similar to that observed at T0. However, at T2 (after 48 hours of recovery) the ratio of glia proliferation was similar to that of control, undamaged discs (Figure S3), suggesting that the signals that promote glia division cease after 48 hours of damage induction.

To evaluate the contribution of cell proliferation on the increased number of glial cells observed in damaged eye discs, we blocked glial proliferation in discs in which apoptosis was induced in the retina region. To this end, we used the *QF/QUAS* system (Potter et al., 2010; Potter and Luo, 2011) in combination with the Gal4 system, enabling us to independently control the expression of QF and Gal4 in the retina and glia cells respectively. To induce genetic ablation, we expressed *QUAS-rpr* in the retina cells under the control of *GMR-QF*. Glia proliferation was simultaneously blocked by expressing a constitutively activated form of the Retinoblastoma Factor (*UAS-rbf^280^*) under the control of *repo-Gal4* driver, which is specifically expressed in glia cells. *rbf^280^* is a mutant form of *rbf* that has four putative cdk phosphorylation sites mutated, and can no longer be regulated by Cyclin D or Cyclin E. The ectopic expression of this factor inhibits dE2F1 activity (Du et al., 1996) and subsequently, cell proliferation. *GMR>rpr tub>Gal80 ^ts^ repo>rbf ^280^* larvae were raised at 25°C until 96 hours after egg laying (AEL) and then shifted to the restrictive temperature to block cell division for 24 hours. After this time, we found that glia cells proliferation was strongly reduced in both damaged and control discs (Figure 3). Accordingly, we detect that the number of glia cells was also sharply reduced and subsequently, the excess of glial cells found in damaged discs was entirely eliminated. However, compared to undamaged discs where proliferation is blocked, in damaged discs we observed more glia cells, suggesting that glia over-migration also contributes to the elevated number of glial cells found in injured eye discs. The induction of *UAS*-*rbf^280^* for a prolonged period of time (72 hours) enhances all the effects caused by the ectopic expression of this factor. Using this later experimental setup, cell proliferation was fully abolished, yet more glial cells were found when compared to undamaged discs in which proliferation was blocked (Figure 3).

**Figure 3.**
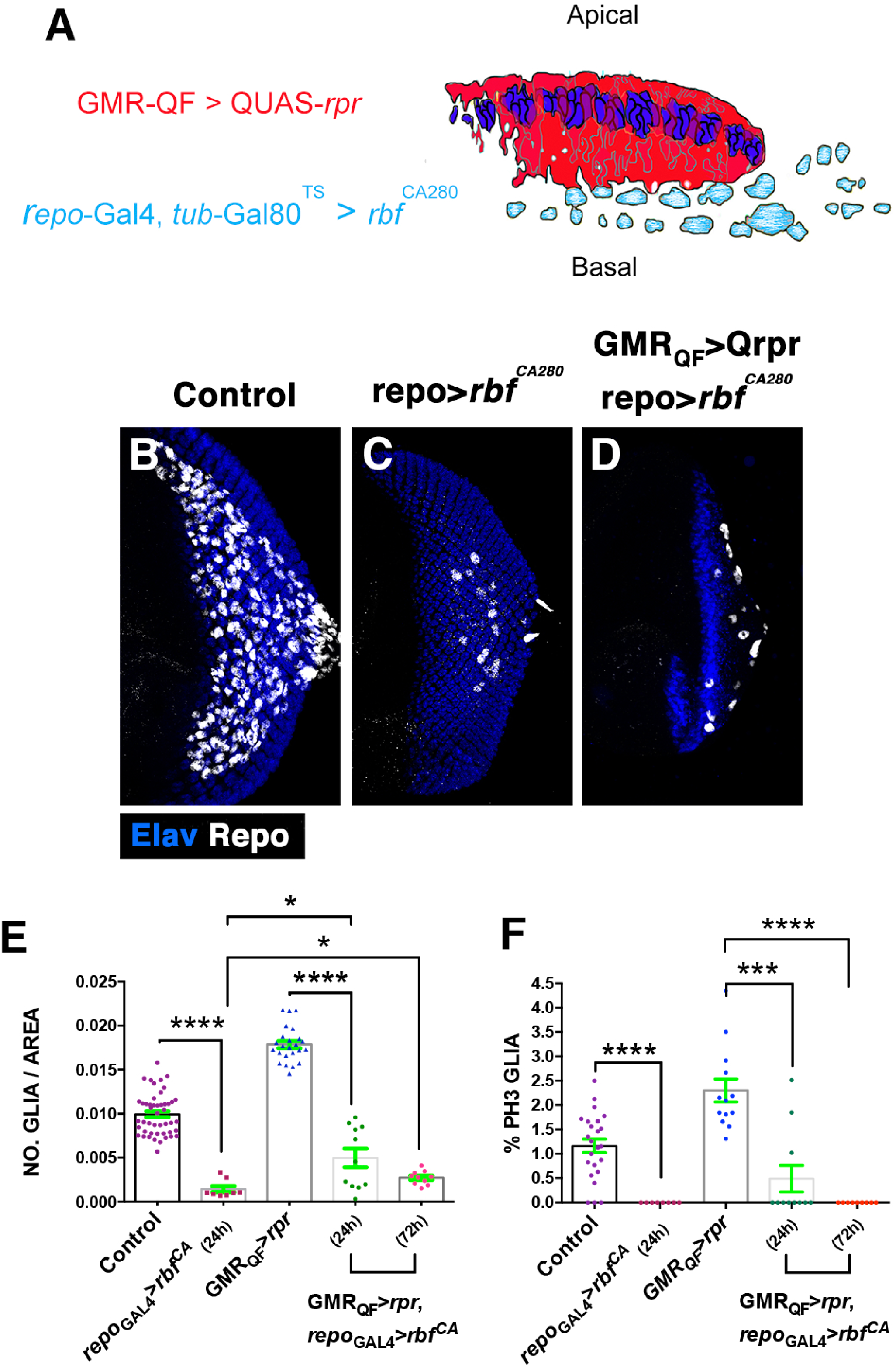
Overexpression of *rbf^CA280^* in glial cells reduces the number of glial cells observed in damaged discs. (A) The Schematic illustration represents a transverse section of an eye disc where the region marked in red (*expression domain of GMR-QF)* corresponds to the area of the discs that has been damaged using *QUAS-rpr GMR-QF*. Glial cells are indicated in light blue. *repo-Gal4* drives the expression of *UAS-rbf^CA280^* under the control of *UAS* specifically in glial cells. (B-D) Third instar eye discs stained with anti-Repo (white) and anti-Elav (blue). Control undamaged *GMR-QF; tub-Gal80^TS^ repo-Gal4* eye disc (B), *tub*-*Gal80^TS^*/+; *repo*-*Gal4*/*UAS-rbf^CA280^ (C) and GMR*-*QF*; *tub*-*Gal80^TS^*/+; *repo*-*Gal4*/*UAS-rbf^CA280^* disc (D). The over-expression of *rbf^CA280^*under the control of *repo-Gal4* reduces the number of glial cells in undamaged (C) and damaged eye discs (D). (E) The graph represent the glial density of discs shown in B-D (Control, *repo_Gal4_-rbf_24hrs_*, *GMR_QF_-rpr*, *GMR_QF_-rpr repo_Gal4_-rbf_24hrs_* and *GMR_QF_-rpr repo_Gal4_-rbf_72hrs_*). (F) The graph shows the % of glial cells in mitosis (Ph3 positive). Error bars represent SEM.

Taken together, our data suggest that after damaging the retina region, signals were generated by apoptotic cells that non-autonomously promote glial proliferation, resulting in an increase in the number of these cells in the eye disc.

### Glial cells produce long cellular projections toward injured region in response to damage

The eye imaginal disc harbours two main glia subtypes, the perineurial and wrapping glia cells. We have examined their behaviour in response to neuronal damage. To this end, we induced genetic ablation in the retina using the QF-QUAs system and visualized glia cells with *UAS-GFP* under the regulation of *Mz97-Gal4* and *c527-Gal4* driver strains, which drive expression in wrapping and perineurial glial cells, respectively.

Damage causes a remarkable change in the behaviour and morphology of wrapping glial cells. In control discs, wrapping glia (WG) send long cellular processes that follow the axons through the optic stalk toward the brain. However, in damaged *GMR-QF; UAS-GFP Mz97-Gal4; QUAS-rpr* discs we did not observe these processes in the optic stalk (Figure S4 compared A-A’ to B-B’). In these discs, the density of wrapping glia, as well as the proportion of these cells in the eye discs was strongly reduced compared to control discs (Figure 4 A-F’ and O-P). In contrast to control discs where WG were always located in the basal layer of the discs, in damaged discs we observed these glial cells in the middle/apical layers of the disc epithelium (Figure 4 D-F’). Moreover, whereas in control discs the anteriormost row of wrapping glia coincided with the anterior leading edge of the glial field, which are close to the morphogenetic furrow, in damaged discs, wrapping glia were located between 5-4 rows of glia cells behind this boundary (Figure S4 compared A with B, and C). Another interesting feature of WG in damaged discs is the increased complexity of their cytoplasmic projections (seen with GFP) compared to wrapping glia in control discs (Figure 4 compared A-C’ with D-F’ and G-G’’). Remarkably, we found that some of these extensions can go over the apical surface of photorreceptors (Figure 4 F-F’). These results were confirmed by performing a time-lapse image analysis to visualise WG cells in controls and damaged discs. In damaged discs we observed that WG cells extend long membranes projection toward the damaged region (Video 1 and Figure 4), whereas we did not observe these projections in control discs (Video 2). In addition, wrapping glial cells in damaged discs contained multiple vesicles with apoptotic neuronal debris inside. Occasionally, we even observed WG enwrapping entire neuronal nuclei stained with the neuronal marker Elav (Figure 4 G-H’’). Another feature of the transformed WGs is that their nuclei were consistently bigger than the nuclei of control WG (Figure 4 Q).

**Figure 4.**
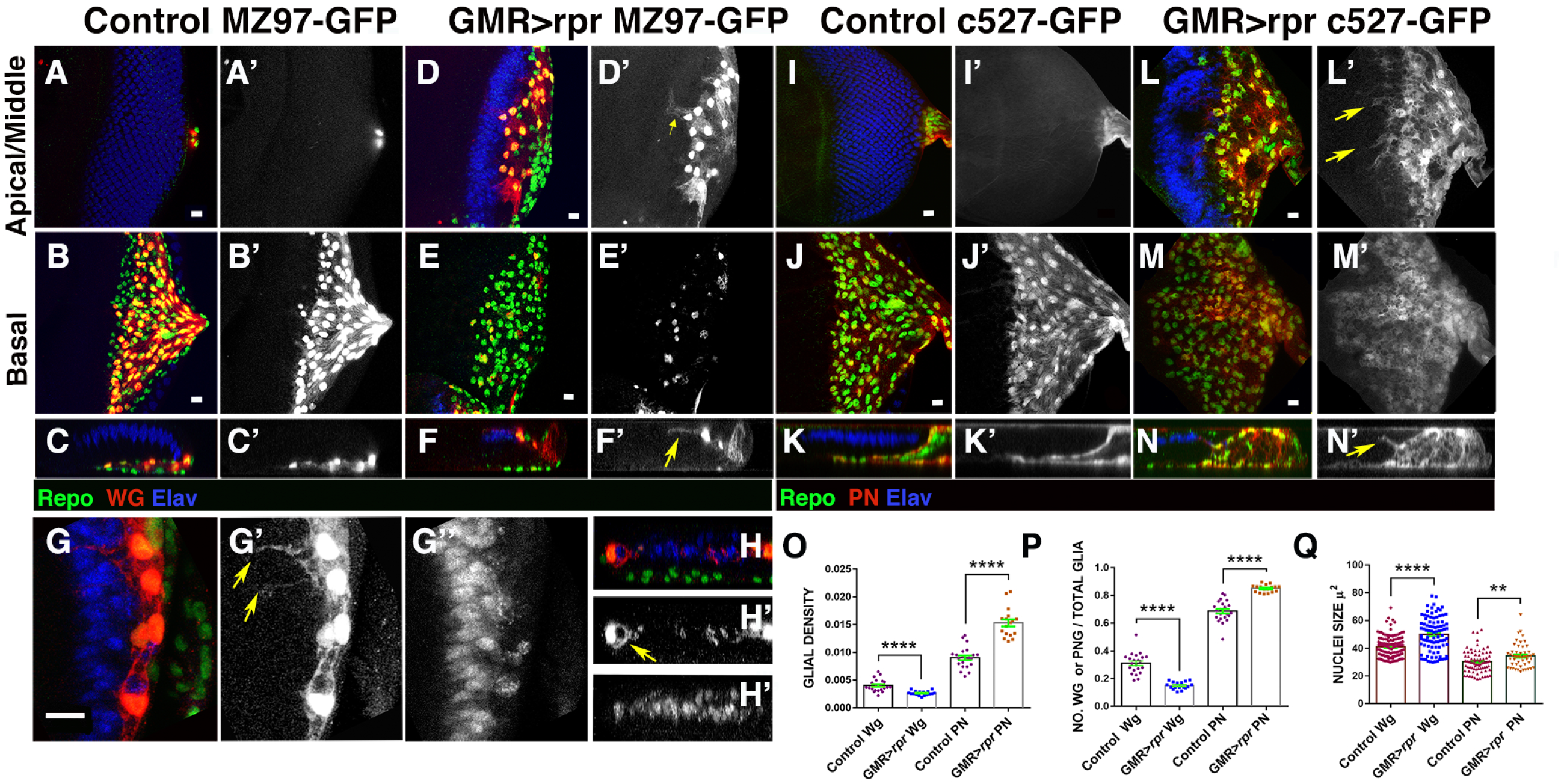
Wrapping and perineurial glial cells change their morphology in response to damage. (A-N) Third instar eye discs stained with anti-Elav (Blue) and anti-Repo (green). (A-H’’) *Mz97-Gal4* driving expression of *UAS-GFP* (in red) reveals wrapping glial cells in control *UAS-GFP Mz97-Gal4; QUAS-rpr* (A-C’) and in damaged *GMR-QF; UAS-GFP Mz97-Gal4; QUAS-rpr* (D-H’’) discs. (I-N) *c527-Gal4* drives expression of *UAS-GFP* (red) in perineurial glial cells in control *UAS-GFP c527-Gal4; QUAS-rpr* (I-K’) and in damaged *GMR-QF; UAS-GFP c527-Gal4; QUAS-rpr* eye discs (L-N). (A-A’, D-D’, I-I’ and L-L’) Apical/middle layers and (B-B’, E-E’, J-J’ and M-M’) Basal layers of the eye disc ephilelium. (C, F, K and N) Cross-sections perpendicular to the furrow of the eye discs shown in A-L’. (A-F’) In control discs we never find wrapping glia cells in the middle or apical layer of the discs (A-C’), however, in damaged discs most wrapping glial cells are located in apical region (D-F’ yellow arrow in F’). These cells extend large processes towards the damaged region (G-G’’, yellow arrows in G’). Some of these projections go over photoreceptors (yellow arrowhead in F’). These processes can enwrap even entire nuclei (yellow arrow in H’), stained with anti Elav (blue). (I-N) In damaged eye discs some perineurial glial cells are located in the apical and middle layers of the disc ephilelium (compared N-N’ with control K-K’) and they send cellular projections toward the damage region (arrows in L’ and N’). (O-Q) Graphs showing: wrapping (Wg) and perineurial (PN) glial density (O), ratio of wrapping (WG) and perineurial (PN) glial cells (n° wrapping or perineurial glial cells/n° Total glia) (P), and nuclei size (Q).

Altogether, our data suggest that damage provokes wrapping glial cells to expand their membrane surface, generating glial processes that engulf and phagocytise cellular debris and apoptotic corpses.

The number of WGs reduces after injury; however, conversely, we found that in damaged discs the density and proportion of perineurial glial cells increased (Figure I-N’, O and P). Moreover, we detected that some perineurial glial (*c527-* expressing cells) have been displaced towards the medial layer of the discs (Figure 4 compared I-K’ with L-N’). In control discs, the cells expressing *c527-Gal4* line are always in the basal layer of the disc (Figure 4 I-K’). Interestingly, we clearly identify that some glial cells expressing *UAS-GFP* under the control of *c527-Gal4* located in middle/apical layers have long cellular processes, similar to those described for WG cells. Moreover, these cells enclose vesicles containing cellular debris labelled with the neuronal marker anti-Elav, suggesting that these perineurial glial cells have phagocytic activity (Figure S4). The average size of the nuclei of these perineurial glial cells is bigger than the nuclei of perineurial glial cells from undamaged discs (Figure Q). Our observations indicate that 27% (SD=±0,07, n=12 discs analysed) of glial cells with large nuclei located in the middle/apical planes correspond to perineurial glia, while the remaining 87% correspond to wrapping glial cells.

### JNK signalling is activated autonomously in damaged regions and non-autonomously in glial cells in response to damage

The c-Jun N-terminal kinase (JNK) pathway plays a fundamental role in regulating many biological processes involved in regeneration, including activation of glial response to damage in the CNS (Kato et al., 2018; Lu et al., 2017; Kato et al., 2011). Moreover, it has been proposed that in developing eye discs the ectopic activation of this pathway in the cells anterior to the morphogenetic furrow is sufficient to induce matrix metalloproteinase 1 (Mmp1) expression and trigger glia over-migration (Tavares et al., 2017).

We have examined the activity of this signalling pathway after damaging the retinal region using the JNK reporter (*TRE-GFP*), as well as a *Lac-Z* insertion in the gene *puckered (puc).* This gene encodes for a phosphatase that controls the activity of JNK-signalling via a negative feedback loop and its expression is a good readout of the activity of the pathway (Martin-Blanco et al., 1998). *TRE-GFP* reporter contains four binding sites for AP-1 (a heterodimeric Jun/Fos transcription factor targeted by JNK) upstream (Chatterjee and Bohmann, 2012).

In third instar control eye discs (*GMR-Gal4 tub-Gal80^ts^ / TRE-GFP*), the *TRE-GFP* reporter is expressed in some photoreceptor cells and unevenly in glial cells (Figure S5). After damaging the discs, the activity of *TRE-GFP* significantly increases in photoreceptor neurons, interomatidial and accessory cells in the damaged region, as well as in some glial cells (Figure S5 I-O’’). We found that the glial cells that express higher levels of *TRE-GFP* are those located in the medial and apical layer of the eye discs epithelium (Figure S5 I-P’’). Similar results were obtained with the *puc-LacZ* line after damage induction. The expression of this reporter increased in damaged regions and in some glial cells that were preferentially located in apical and middle layers of the discs (Figure S6). To further characterize the activity of JNK signalling in glial cells in response to damage, we analysed the genomic region of *puc* to identify a possible non-coding regulatory region that could be used specifically as a reporter of this signal in glia cells. To this end, we took advantage of the published open chromatin profile of eye discs during tumour development. We selected two different regions (*puc-1* and *puc-2*, see Figure S7) and cloned them into a *lacZ* reporter gene construct. Interestingly, *puc-2* is expressed in glia cells in the eye disc, and a sub-fragment of this element (named *puc-2B*) reproduced this pattern of expression (Figure S7 and Figure 5). The proportion of glial cells that express this reporter is similar in both damaged and undamaged control discs (Figure 5 I). However, damaged lead to up-regulation of *puc2B-LacZ* levels in some glial cells preferentially located in the middle/apical layers of the eye epithelium. This can be most clearly seen in a projection of several transverse sections generated from different damaged discs (Figure 5 H-H’’ compared to control discs D-D’’). Consequently, the proportion of glial cells with elevated levels of *puc2B-Lacz* in these layers of the discs was higher than in the basal layer (Figure 5 J).

**Figure 5.**
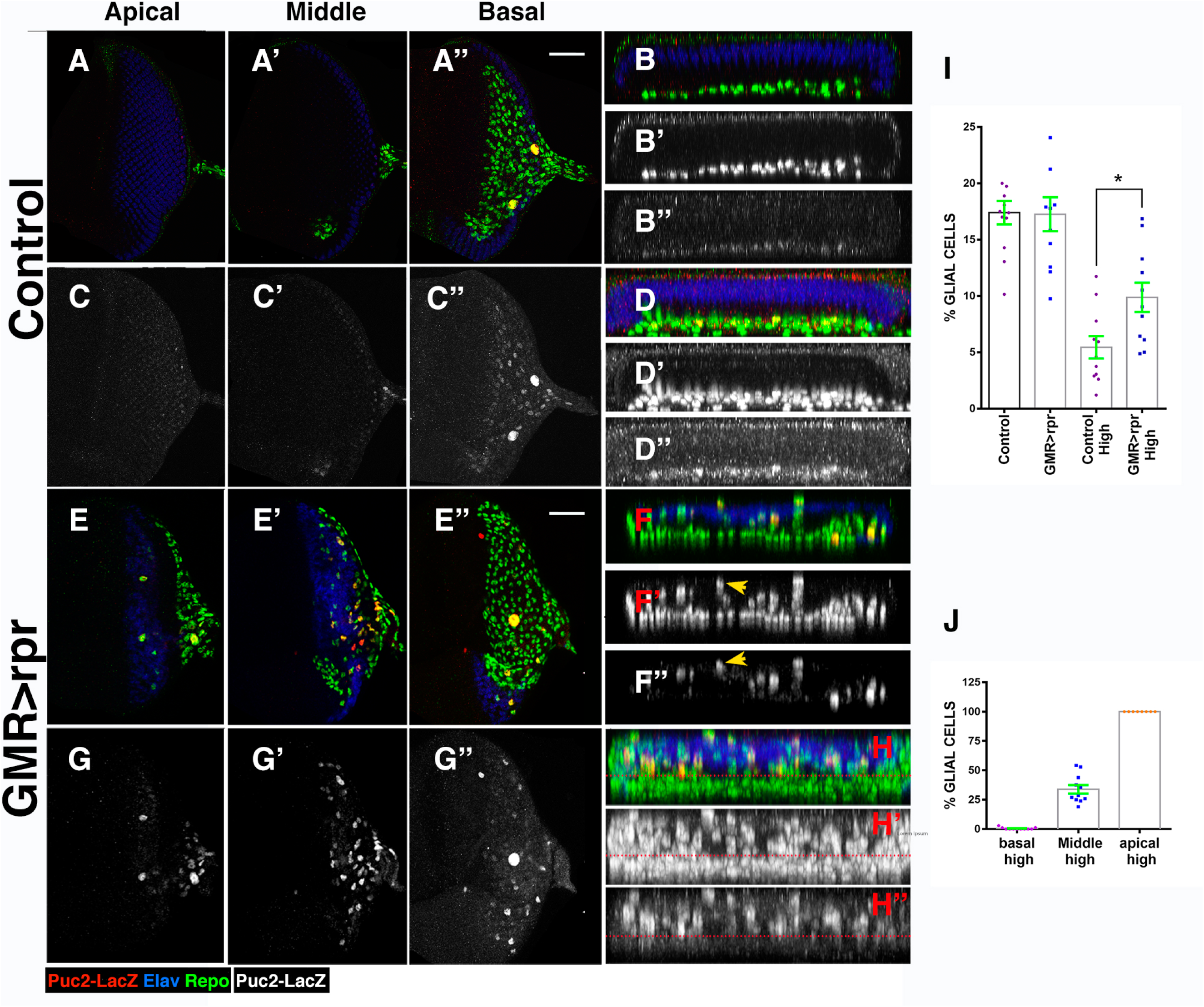
JNK signalling is activated in response to damage. (A-H’’) Third instar eye discs stained with anti-Elav (blue), anti-Repo (green) and anti-B-galactosidase (red) to reveal the activity of the *puc2b-LacZ* reporter, in control *(GMR-Gal4 tub-Gal80^ts^/+; puc2b-LacZ/+*) (A-D’’) and damaged *UAS-rpr/+; GMR-Gal4 tub-Gal80^ts^ /+; puc2b-LacZ/+* discs (E-H’’). Apical (A, C, E, and G) Middle (A’, C’ E’, and G’) and basal (A’’, C’’ E’’ and G’’) layers of control (A-A’’ and C-C’’), and damaged (E-E’’ and G-G’’) eye discs. (B-B’’ and F-F’’) Y–Z projections show two cross-sections parallel to the furrow in control (B-B’’) and damaged (F-F’’) discs. (D-D’’ and H-H’’) Projection of tranversal sections generated from 5 different control discs (D-D’’) and damaged discs (H-H’). Glial cell with higher levels of *puc2b-LacZ* are preferentially located in apical and middle layers of the discs (yellow arrows in F’ and F’’). In the plot shown in H-H’’ note that most glial cells expressing high levels of the reporter are located above the red dotted line. (I) Graph shows % of glial cells expressing *puc2-lacZ* (n° cell expressing *puc2b-LacZ*/Total n° of Glia) in control (control) and damaged *UAS-rpr/+; GMR-Gal4 tub-Gal80^ts^/+; puc2b-LacZ/+* (*GMR<rpr*), as well as the % of glial cells expressing this reporter at high levels (n° cell expressing *puc2b-LacZ* at high levels/Total n° of Glia) (see method). (J) Graph shows % of glial cells expressing *puc2b-LacZ* at high levels (high) in basal, middle and apical layers of damaged eye discs.

### JNK signal is required for promoting glial migration during GRR

The results described so far indicate that in response to damage, JNK signalling was activated in both the damaged region, as well as in glial cells. Therefore, it is plausible that its function may be promoting some aspect of glial response in damaged eye discs. To complement the expression analysis, we examined the glial response after modifying JNK signalling activity in both the damaged region and glial cells.

Firstly, we analysed the autonomous requirement of JNK signalling in the damaged region by depleting its function in the injured domain. To that end, we co-expressed a dominant negative form of the kinase Basket (*UAS-bsk^DN^*) and *UAS-rpr* under the control of a *GMR-Gal4* line. Basket is involved in the transduction pathway of JNK; phosphorylating c-Jun and c-Fos transcription factors (Sluss et al., 1996; Weston and Davis, 2007). *UAS-bsk^DN^/UAS-rpr; GMR-Gal4 tub-Gal80^ts^* larvae were transferred for 4 days at restrictive temperature to induce genetic ablation while over-expressing *UAS-bsk^DN^*. We detected a moderate but reproducible and statistically significant reduction in the density of glial cells present in the eye discs of these larvae (Figure 6). The density of apoptotic cells in these discs was similar when compared to discs over-expressing only *UAS-rpr*, suggesting that this suppressive effect was not due to attenuation of the damaged induced by the over-expression of *rpr* (data not shown). Interestingly, we found that there were no statistically significant differences in glial cell proliferation between these discs and control damaged discs (Figure 6 I). The over-expression of *UAS-bsk^DN^* alone (*bsk^DN^; GMR-Gal4 tub-Gal80^ts^*) did not cause any detectable change compared to control discs. These results suggest that the function of the JNK pathway is required in apoptotic cells to promote glial response.

**Figure 6.**
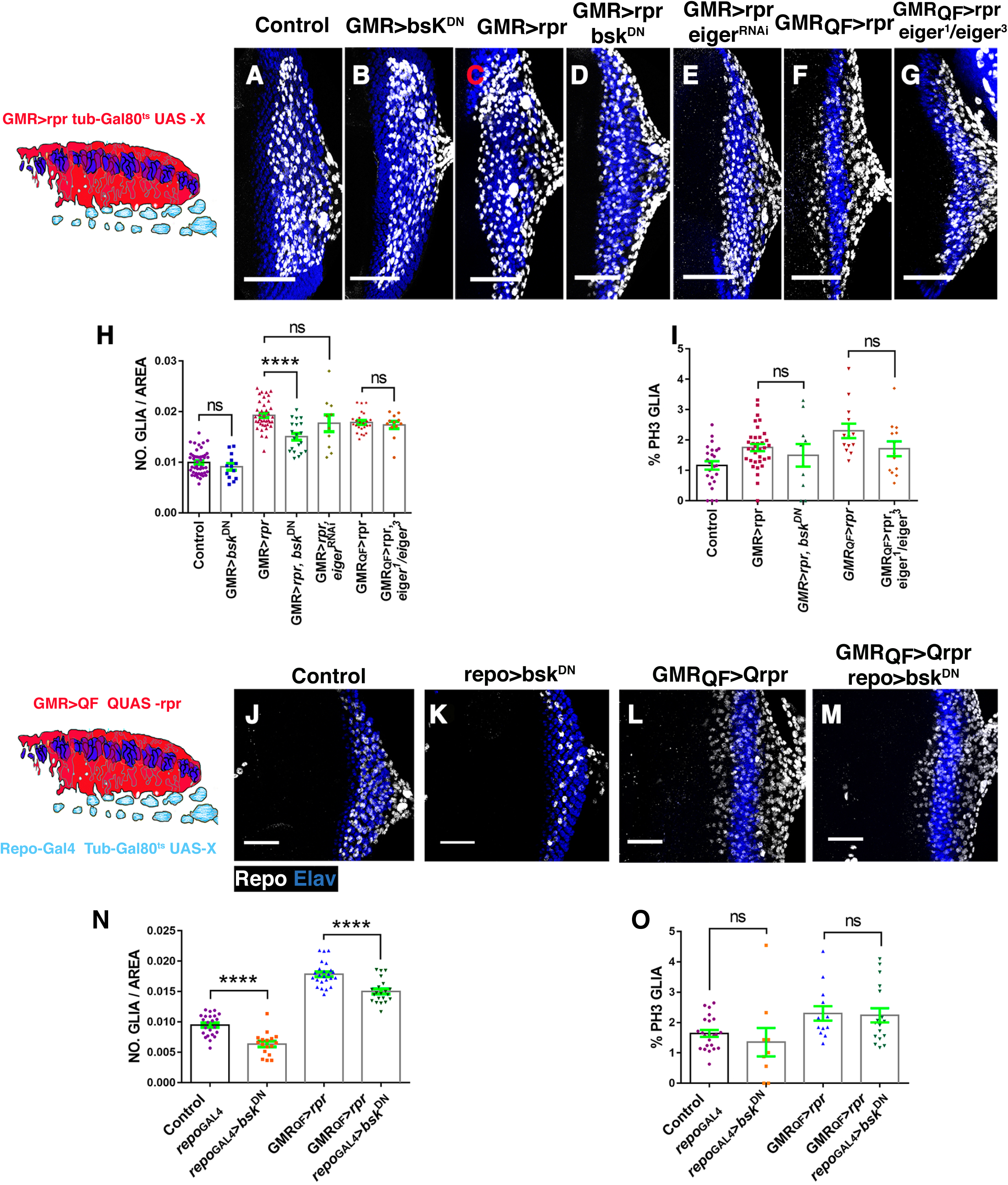
Reduction of JNK activity prevents the accumulation of glial cells produced upon damage. (A-G’’ and J-M) Projections of confocal images of third instar eye discs stained with anti-Elav (blue) and anti-Repo (white). The Schematic illustration on the left represents a transverse section of an eye disc where the region marked in red (*expression domain of GMR-Gal4)* corresponds to the area of the discs that has been damaged at the same time that JNK was depleted. (A-E) Non-autonomous effects on glial cells caused by the down-regulation of JNK signalling in the damaged region. (A) Control undamaged *GMR-Gal4 tub-Gal80^ts^*/+ disc. (B) *UAS-bsk^DN^; GMR-Gal4 tub-Gal80^ts^* eye. (C) Damaged *UAS-rpr/+; GMR-Gal4 tub-Gal80^ts^*dics. (D) *UAS-rpr/UAS-bsk^DN^; GMR-Gal4 tub-Gal80^ts^* damaged discs. (E) *UAS-rpr/; GMR-Gal4 tub-Gal80^ts^/ UAS-eiger ^RNAi^*. (F) *GMR-QF; QUAS-rpr* eye discs damaged. (G) Damaged *eiger* mutant disc *GMR-QF; eiger^1^/eiger^3^; QUAS-rpr*. All larvae were subjected to the same temperature shifts (see method) except larvae of the discs shown in F and G that were raised at 25 °C throughout the development. (H-I) Graphs show glial cell density (H) and % of mitotic glia cells of discs shown in A-G. (J-M) Effects of the down-regulation of JNK in glial cells of damaging eye discs. In the schematic illustration on the left is indicated in red the region that has been damaged by over-expressing *QUAS-rpr* under the control of *GMR-QF* and in light blue glial cells. (J) Control undamaged *tub-Gal80^ts^ repo-Gal4* disc. (K) *UAS-bs^DN^* /+; *repo-Gal4/+* eye discs. (L) Damaged *GMR-QF; tub-Gal80^ts^*; *repo-Gal4 QUAS-rpr* eye disc. (M) *GMR-QF UAS-bsk^DN^; tub-Gal80^ts^ repo-Gal4 QUAS-rpr discs.* (N-O) Graphs show glial cell density (N) and % of mitotic glia cells (O) of discs shown in J-M.

We next examined whether the JNK pathway was also required in glial cells. To do this, we combined the *QF/QUAS* system to induce damage and the *Gal4/UAS* to block JNK pathway signalling in glia under the control of *repo-Gal4* (Figure 6). In *GMR-QF>rpr repo-bsk^DN^* discs, we observed a moderate and statistically significant reduction in the density of glia cells. The proliferation rate of glial cells was compared with control damaged eye discs from larvae that were subjected to the same temperature shifts (Figure 6 N and O). The over-expression of *UAS-bsk^DN^* in glial cells under the same experimental condition, reduced the density of glial cells present in the eye discs, but did not affect glial cell proliferation (Figure 6 N and O).

These results suggest that the JNK pathway is required in both the damaged region and in glial cells to increase glial cell number in response to damage. However, JNK signalling appears to be dispensable for inducing glial proliferation, suggesting that this pathway is instead necessary to promote glial cell motility. In an attempt to better understand the function of JNK signalling in regulating glia behaviour during development, we examined the pattern of proliferation and motility of glial cells in an *hemipterous* (*hep*) mutant background. This gene encodes a serine/threonine protein kinase involved in the transduction of JNK signalling. In *hep^r75^* mutant eye discs, the total number of glia in the eye disc was significantly reduced, but again without affecting glial proliferation. Both the % of dividing glial cells, as well as EDU incorporation were similar to that seen in control eye discs (Figure S8). To obtain an overview of glial motility in *hep^r75^* mutant discs, we performed a time-lapse image analysis where glial cells were visualised with *UAS-GFP* driven by *repo-Gal4.* Glial cells migrate by generating short cellular projections spreading toward the leading edge. We observed these projections in both control and *hep^r75^* mutant discs (Figure S9 and video 3 and 4). However, in control discs glial cells move forward steadily, whereas in *hep^r75^* mutant discs, most glial cells remained in the same position or even moved backwards, with only few cells moving slightly forward (Figure S9 and Video 3 and 4). Therefore, the average speed and average displacement of glial cell in *hep^r75^* mutant discs was significantly reduced compared with control discs (Figure S9).

Altogether, our data suggest that JNK signalling is required in both the damaged region, as well as in glial cells, to promote their motility.

Our results also imply that apoptotic cells produce a signal that non-autonomously activates JNK signalling in glial cells. The tumor necrosis factor (TNF) ligand termed Eiger is an obvious candidate for being this effector; leading us to then examine the effect of the loss of *eiger* (the *Drosophila* TNF homolog), on glial response. We first blocked *eiger* activity in the damaged region by co-expressing *UAS-eiger^RNAi^* and *UAS-rpr* under the control of *GMR-Gal4*. The density and proliferation of glial cells in these discs was similar to that found in control damaged discs (Figure 6). We obtained the same result when analysing these parameters in *eiger^1^/eiger^3^* mutant discs eye discs that have been damaged (*QMR-QF; eiger^1^/ eiger^3^; QUAS-rpr*) (Figure 6). In *eiger^1^/eiger^3^* mutant discs, glial cells were not affected (Figure S8). These results suggest that Eiger was not the signal emitted by apoptotic cells to activate JNK in glial cells. This is supported by our observation that the overexpression of *UAS-eiger* under the control of *GMR-Gal4* only leads to the up-regulation of *TRE-GFP* reporter autonomously, in the GMR domain, but not in glial cells (data not shown).

To investigate whether the activation of JNK signalling was sufficient to increase the number of glial cells in the eye disc, we examined the consequence of transiently expressing a constitutively activated form of the JNK-kinase Hemipterous (*hep^CA^*). The ectopic expression of *UAS-hep^CA^* under the control of *GMR-Gal4* causes a moderate but re-producible and statistically significant increase in glial density compared to control discs (Figure S10). As previously reported by others (Ma et al., 2012; Wang et al., 2018), we found that in discs over-expressing *GMR-Gal4 UAS-hep^CA^,* apoptosis strongly increases (data not shown). Therefore, it is possible that the augmented number of glial cells found in these discs might be a consequence of cell death induction. To further examine this idea, we analysed the effects of co-expressing *UAS-hep^CA^* and *UAS-RHG microRNA* (miRNA). This transgene generates miRNAs that simultaneously inhibit the function of the pro-apoptotic genes, *reaper*, *hid*, and *grim* (Siegrist et al., 2010). In eye discs co-expressing *UAS-hep^CA^* and *UAS-RHG* under the control of *GMR-Gal4* (*GMR-Gal4/ tub-GAL80^ts^ UAS-RHG UAS-hep^CA^*) the excess of glial cells found upon over-expression of *UAS-hep^CA^* was totally suppressed (Figure S10), suggesting that the augmented number of glial cells caused by *UAS-hep^CA^* was due to the induction of apoptosis.

We next examined whether the activation of *UAS-hep^CA^* in glial cells could increase their number within eye discs. *tub-Gal80^ts^; repo-Gal4/UAS-hep^CA^* larvae were shifted to 29°C for 24, 48 and 72 hours to induce *UAS-hep^CA^* activation. Larvae transferred for more than 24 hours at restrictive temperature could not be recovered due to lethality. Therefore, we could only analyse and quantify the effects on eye discs from larvae in which *UAS-hep^CA^* was over-expressed for 24 hours. The number and localization of glia cells presented in these discs were comparable to those observed in control discs, suggesting that the activation of JNK signalling in glial cells was not sufficient for inducing their over-migration into the eye disc (Figure S10).

### *hh* and *dpp* signalling are ectopically activated in damaged eye discs

The diffusible factors Decapentaplegic (Dpp) and Hedgehog (Hh) play an important role on the development of eye discs. The newly differentiated photoreceptors behind the MF express Hh that can diffuse and act on cells anterior to the MF (Heberlein et al., 1995). Hh is known to induce the expression of the BMP2/4 type morphogen *decapentaplegic* (*dpp*) ahead of the MF. This diffusible factor functions redundantly with Hh in MF progression (Curtiss and Mlodzik, 2000; Greenwood and Struhl, 1999; Heberlein et al., 1993; Wiersdorff et al., 1996). Both Hh and Dpp were also suggested to regulate glial cell development in the eye disc. Initially, Hh prevents precocious glial migration (Hummel et al., 2002); in later stages, Hh stimulates glial motility, whereas Dpp induces glial proliferation (Rangarajan et al., 2001). Thus, we wondered if they might be involved in the activation of glial response to eye disc damage. To study this, we first examined the expression pattern of a *dpp-LacZ* line in discs expressing *UAS-rpr* under the control of *GMR-Gal4*.

In third instar control discs, *dpp-lacZ* is expressed along the furrow and is abruptly downregulated posterior to the furrow. However, in damaged eye discs, we observed ectopic patches of *dpp-lacZ* in the region posterior to the MF (Figure 7). We further analysed the response of this signalling pathway after damage by examining its activation via the staining of a phosphorylated form of the transcriptional effector Mad (pMad). This transcription factor is phosphorylated when *dpp* signal is active and therefore can be used a read out of this signalling pathway (Affolter et al., 2001). In control third instar eye discs, pMad staining was restricted to a band of cells adjacent to the MF, and its expression decreased posteriorly (Figure S11). However, in damaged discs pMad is expressed at high levels in most of the cells posterior to the MF. Most notably, we observed that pMad staining increases in some glia cells (Figure S11).

**Figure 7.**
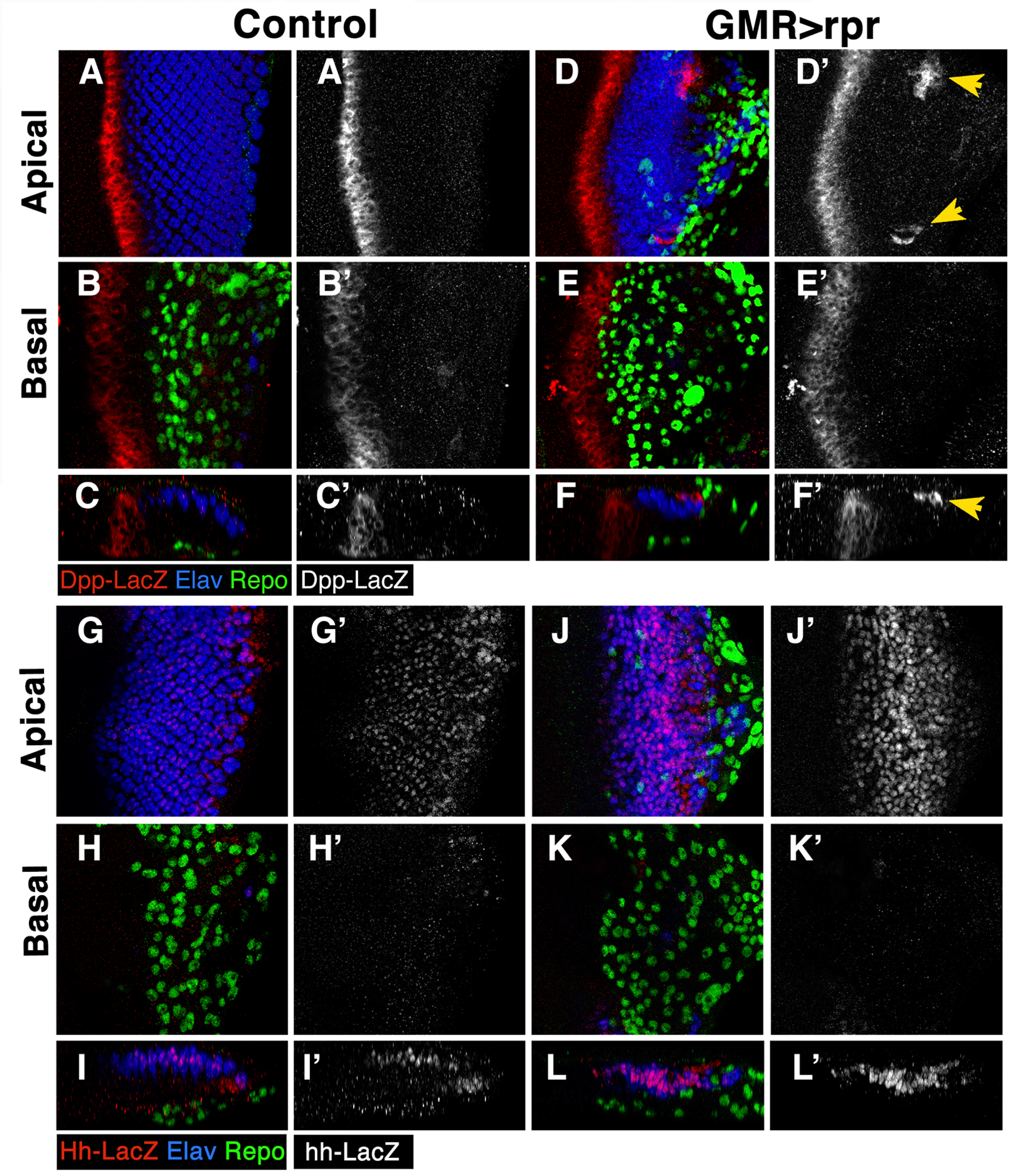
*dpp* and *hh* expression increase in damaged discs. (A-L’) Third instar eye discs stained with anti-Elav (Blue), anti-Repo (green) and anti-ß-Galactosidase to detect the *dpp-lacZ* expression (in red A-C, D-F, and in grey A’-C’, D’-F’), or *hh-LacZ* (in red G-I, J-L and grey G’-I’, J’-L’). (A-A’, D-D’, G-G’ and J-J’) Apical/middle layers and (B-B’, E-E’, H-H’ and K-K’) Basal layers of the eye disc ephilelium. (C-C’, F-F’, I-I’ and L-L’) Cross-sections perpendicular to the furrow of the eye discs shown above. (A-C’) In control discs *dpp-LacZ* is expressed in a band of cells along the furrow. However, in injured *UAS-rpr; dpp-LacZ/GMR-Gal4 tub-Gal80^ts^* discs we observed patches of cells expressing this reporter in the apical layer of the disc (yellow arrowheads in D-D’, and F-F’), compared D-F’ to control discs A-C’. (G-L’) In control discs *hh-LacZ* is expressed in the photoreceptor and accessory cells (G-I’). However, in injured discs the reporter signal increased in the photoreceptor and accessory cells surrounding the wound (J-L’ compared to control discs G-I’).

Next, we analysed *hh* expression upon damage induction by examining the activity of an *hh-lacZ* enhancer trap line in *UAS-rpr; GMR-Gal4 tub-Gal80^ts^* eye discs. In control discs, this reporter was expressed at low levels in photoreceptors (Figure 7). However, in damaged discs the activity of this reporter was up-regulated in photoreceptor cells, as well as in some interomatidial cells (Figure 7). This is consistent with previous data that has shown that the ectopic expression of the pro-apoptotic gene *hid* in the eye disc under control of *GMR-Gal4* increases *hh* signalling, whose function is required to promote compensatory proliferation (Fan and Bergmann, 2008).

We next examined whether increased levels of *hh* expression in retinal cells might activate *hh* signalling in subretinal glia by analysing the expression of Patch, a known target of this signalling pathway. Consistent with our previous results, we observed that the levels of Patch were abnormally high in cells posterior to the morphogenetic furrow, as well as in subretinal glial cells (Figure S11).

These data suggest that apically located photoreceptor neurons and interommatidial cells up-regulate *dpp* and *Hh* in response to damage. These factors in turn, non-autonomously would activate these two signalling pathways in glial cells.

### *dpp* and *hh* signalling are necessary to promote Glial response

Our observations indicate that in response to damage, both *dpp* and *hh* signalling were activated in the injured region, as well as in subretinal glial cells; leading us to consider if these signals might be involved in stimulating glial response to damage. To assess this, we first examined whether the depletion of *dpp* and *hh* alter the density and/or proliferation of glial cells observed in damaged discs. To this end, we performed RNAi-mediated knockdowns by overexpressing *UAS-hh ^RNAi^* and*/*or a number of different *UAS*-*dpp^RNAi^* lines under the control of *GMR-Gal4* whilst simultaneously inducing damage using *UAS-rpr*. None of the *UAS-dpp^RNAi^* lines employed in our assay significantly modified the effects on glia number observed in damaged discs (Figure 8 and Figure S12). We obtained similar results when the function of *hh* signalling was knocked down upon induction of damage using the same experimental setup (Figure 8).

**Figure 8.**
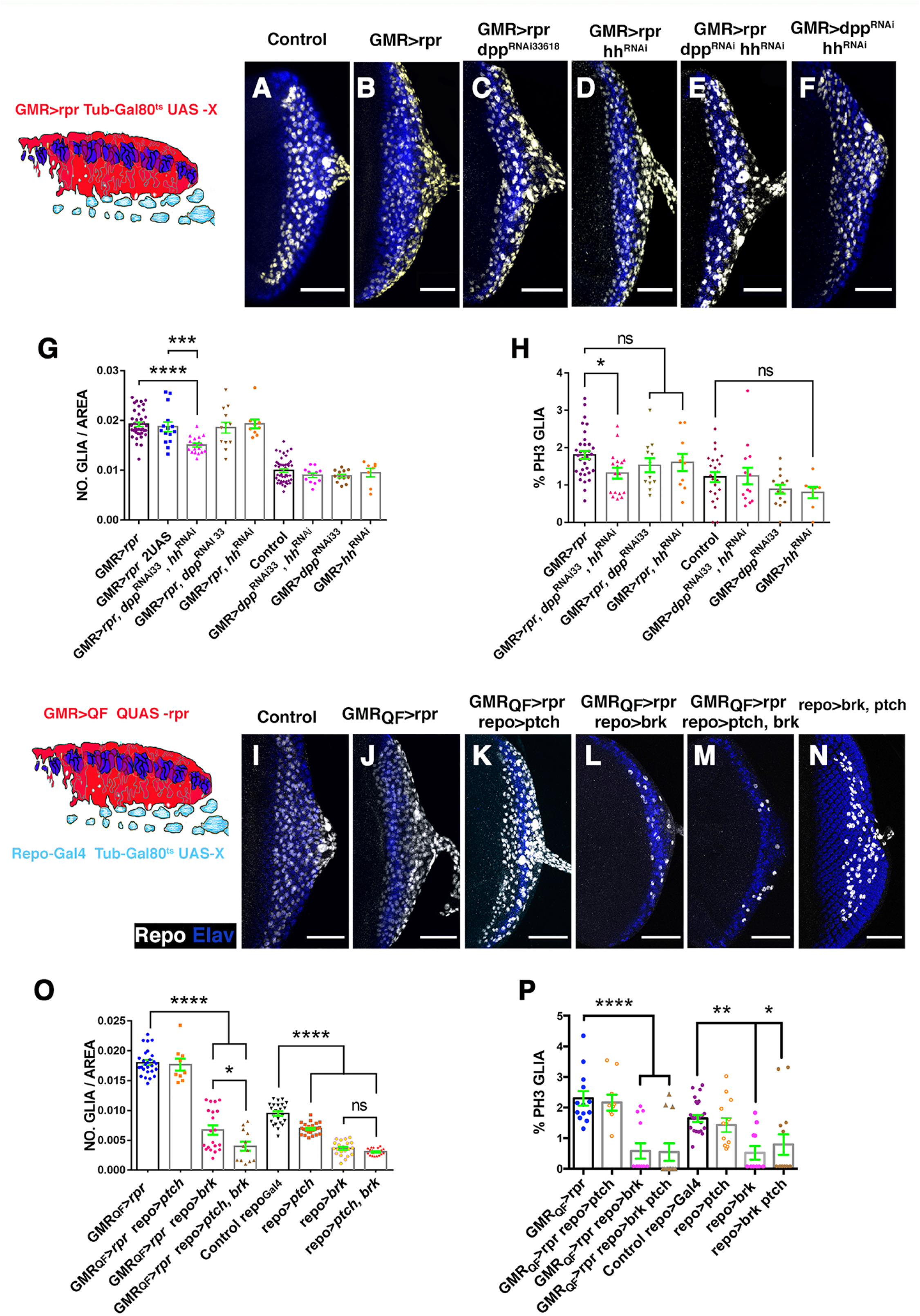
The down-regulation of *dpp* and *hh* signalling reduce glial response. (A-F and I-N) Third instar eye discs stained with anti-Elav (blue) and anti-Repo (white). (A-F) Effects caused by the down-regulation of *dpp* and *hh* signalling in the damaged region. The Schematic illustration on the left represents a transverse section of an eye disc where the region marked in red (*expression domain of GMR-Gal4)* corresponds to the area of the discs that has been damaged at the same time that *dpp* and/or *hh* signalling were depleted. (A) Control undamaged *GMR-Gal4 tub-Gal80^ts^*/+ disc. (B) *UAS-rpr; GMR-Gal4 tub-Gal80^ts^* damaged eye disc. (C) Damaged *UAS-rpr/+; GMR-Gal4 tub-Gal80^ts^; UAS-dpp^RNAi33^* (BDSC33618). (D) *UAS-rpr/+; GMR-Gal4 tub-Gal80^ts^/ UAS-hh^RNAi^* damaged discs. (E) Damaged *UAS-rpr/; GMR-Gal4 tub-Gal80^ts^/ UAS-hh^RNAi^*; *UAS-dpp^RNA33i^*. (F) *GMR-Gal4 tub-Gal80^ts^/ UAS-hh^RNAi^*; *UAS-dpp^RNAi^* eye discs. (G-H) Graphs show glial cell density (G) and % of mitotic glia cells of discs shown in A-F. (I-N) Effects of the down-regulation of *dpp* and *hh* signalling in glial cells in damaged discs. In the schematic illustration on the left is indicated in red the region that has been damaged (*GMR-QF*; *QUAS-rpr*) and in light blue glial cells. (I) Control undamaged *tub-Gal80^ts^; repo-Gal4* disc. (J) *GMR-QF; tub-Gal80^ts^*; *repo-Gal4 QUAS-rpr* eye disc. (K) *GMR-QF; tub-Gal80^ts^/UAS-patch*; *repo-Gal4 QUAS-rpr.* (L) *GMR-QF; tub-Gal80^ts^*; *repo-Gal4 QUAS-rpr/UAS-brk.* (M) *GMR-QF; tub-Gal80^ts^/UAS-patch*; *repo-Gal4 QUAS-rpr/UAS-brk.* (N) *tub-Gal80^ts^/UAS-patch*; *repo-Gal4 /UAS-brk.* (O-P) Graphs show glial cell density (O) and % of mitotic glia cells (P) of discs shown in I-N. Scale bar 50 μm.

Next, we tested whether the simultaneous depletion of *dpp* and *hh* in injured retinal affect glial response by co-expressing *UAS-dpp^RNAi^*, *UAS-hh^RNAi^* and *UAS-rpr* under the control of *GMR-Gal4*. We found that loss of *dpp* and *hh* together, impairs the accumulation of glia in damaged eye disc. The number of glial cells observed in these discs was significantly lower than numbers observed in control damaged discs (Figure 8). Accordingly, we found that compared to control damaged discs, glial cell proliferation was reduced (Figure 8). The down-regulation of *dpp* and *hh* under the control of *GMR-Gal4* using the same experimental conditions neither affect the density or the division of glial cells in undamaging conditions (Figure 8 G and H).

To confirm that this effect was not due to a titration of GAL4 activity caused by introducing an additional transgene, we used damaged eye discs that have two *UAS* reporters as a control (see method) (Figure 8G). These larvae were subjected to the same temperature shifts. In this experimental setup, the effects on glial density were similar to those found in control damaged discs with only one *UAS* construct (Figure 8G).

These results are consistent with a model in which Dpp and Hh are secreted from the damaged retinal region to promote glial response. We therefore examined the consequence of inducing damage in the retinal region and simultaneously blocking *dpp* and/or *hh* pathways in glial cells. Once again, we combined *QF/QUAS* and *Gal4/UAS* system. To block *dpp* signalling in glia we over-expressed *brinker* (*brk*) or *daughters against dpp* (*dad*), under the control of *repo-Gal4. brinker* encodes for a sequence-specific transcriptional repressor that negatively regulates Dpp-dependent genes, whereas *dad* encodes the inhibitory SMAD in the BMP/Dpp pathway, and subsequently downregulates dpp signalling activity (Tsuneizumi et al., 1997).

The over-expression of *UAS-brk* or *UAS-dad* in the glial cells of damaged discs was sufficient to prevent the accumulation of glia induced upon damage. Thus, the glia density in these discs was significantly lower than in control damaged discs (Figure 8 and Figure S12). Also, glia proliferation decreased in damaged discs expressing *UAS-brk or UAS-dad* (Figure 8). The over-expression of these genes under the control of *repo-Gal4* in undamaged discs using the same experimental setup also caused a significant reduction in the number of glial cells, as well as, in their rate of proliferation (Figure 8 and Figure S12). These results imply that *dpp* signalling is necessary for increasing the number of glial cells in response to damage.

We also analysed the effects of blocking *hh* signalling in glial cells from damaged eye discs. To that end, we silenced the function of *cubitus interruptus (ci)* in glial cells by over-expressing an *UAS-ci^RNAi^* under the control of *repo-Gal4*. Alternatively, we over-expressed the negative regulator of the *hh* pathway *patch* under the control of *repo-Gal4*. We observed that in damaged discs the down-regulation of *hh* signalling did not cause any significantly change in the number of glial cells compared to control damaged eye discs (Figure 8 and Figure S12).

In line with our previous observations, we found that in damaged eye discs where both *hh* and *dpp* pathways were simultaneously depleted in glial cells by co-expressing *UAS-dad* and *UAS-ci^RNAi^* or *UAS-brk* and *UAS-patch* under the control of *repo-Gal4,* the number of glial cells was significantly reduced compared with discs in which only *dpp* signalling was depleted (Figure 8 and Figure S12). Interestingly, the proliferative ratio of the glial cells in these discs was similar to that found when only *dpp* signalling was reduced, suggesting that *dpp* plays a more important role in the control of glial proliferation than *hh* signalling.

We do not fully understand why the depletion of *dpp* signalling in glial cells was sufficient to prevent glial accumulation in damaged discs, whereas the silencing of this signalling alone in the damaged region did not affect glial response. A possible explanation is that, as it has been previously reported during eye discs development, *hh* and *dpp* have a partially redundant function (Curtiss and Mlodzik, 2000). Alternatively, the silencing of *dpp* signalling using RNAi may be weak and therefore, only in a sensitised background (i.e. in our assay with the down-regulation of hh signalling), will the down-regulation of *dpp* signalling cause an observable effect in glial response.

Altogether, our data suggest that in response to damage Dpp and Hh are produced and secreted by apoptotic cells in the retinal region, that in turn activate these signalling pathways in glial cells to promote its proliferation and motility. Consequently, we expect that the ectopic activation of these signalling pathways in the glial cells should promote glial proliferation and motility leading to an excess of glia in the eye disc. To this end, we over-expressed a constitutively activated version of the receptor Thickvein (*tkv^QD^*) and *Interference hedgehog (Ihog)* under the control of *repo-Gal4* (Das et al., 1998; Nellen et al., 1994; Pignoni and Zipursky, 1997). Tkv is one of the receptors of Dpp, whereas, *ihog* encodes for a transmembrane protein that is essential for Hh pathway activation (Camp et al., 2010). We observed that numbers of glial cells in eye disc over-expressing *UAS-ihog* under the control of *repo-Gal4* was similar to that of control discs (Figure 9). We confirmed that the over-expression of *UAS-ihog* was sufficient to up-regulate *hh* signalling by observing high Patch expression levels in these discs (data not shown). In contrast, the over-expression of *UAS-tkv^QD^* (*repo-Gal4 UAS-tkv^QD^)* caused a mild increase in the density and proliferation of subretinal glia. These effects were strongly enhanced when we co-overexpressed *UAS*-*tkv^QD^* and *UAS-ihog* under *repo-Gal4* regulation (Figure 9). Interestingly, we also observed that the co-activation of both signalling pathways in glia cells promotes the over-migration of these cells. Whereas in most discs over-expressing either *UAS-tkv^QD^* or *UAS-ihog,* the anterior border of glial migration lies 2-4 rows of ommatidia posterior to the morphogenetic furrow, a high percentage of discs over-expressing simultaneously both *UAS-tkv^QD^* and *UAS-ihog* shown glial cells overcoming that border (Figure 9 D-H).

**Figure 9.**
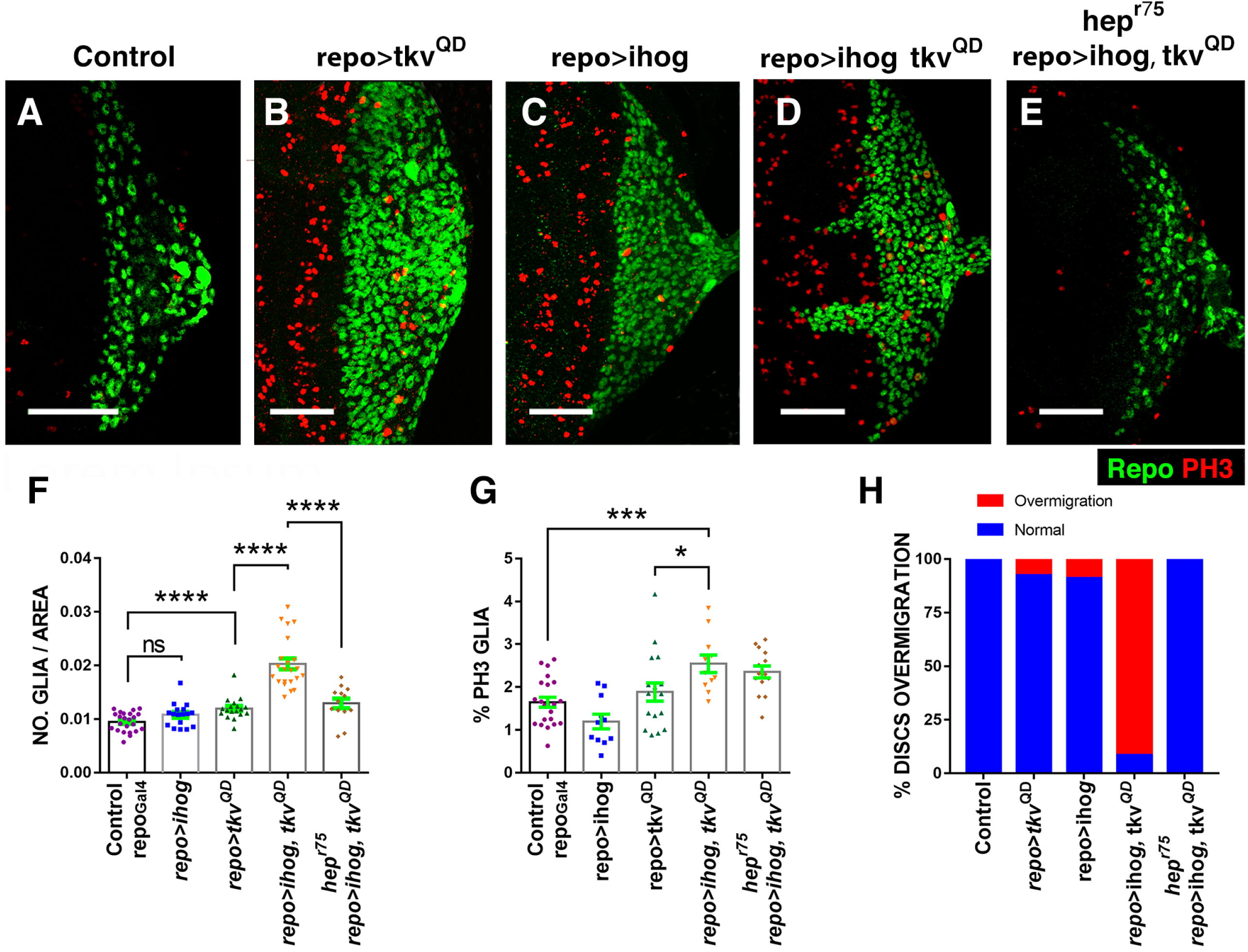
Overexpression of *ihog* and *tkv* in glial cells induces over-migration and proliferation. (A-E) Third instar eye discs stained with anti-PH3 (red) and anti-Repo (green). Basal layers of the eye disc epithelium. Control *tub-Gal80^TS^ repo-Gal4* eye disc (A), *tub-Gal80^TS^*/+; *repo-Gal4/UAS-tkv^QD^* (B), *tub-Gal80^TS^*/*UAS-ihog; repo-Gal4*/+ (C), *tub-Gal80^TS^*/*UAS-ihog*; *repo-Gal4/UAS-tkv^QD^* (D) and *hep^r75^*, *tub-Gal80^TS^*/*UAS-ihog; repo-Gal4/UAS-tkv^QD^* (D). The over-expression of *tkv^QD^* under the control of *repo-Gal4* increases the number of glial cells (B). The total number of glia in eye discs *repo-Gal4 UAS-ihog* appears unchanged with respect to the control (C). The co-overexpression of *UAS-tkv^QD^* and *UAS-ihog* under *repo-Gal4* increases the density and proliferation of subretinal glial cells and promotes the over-migration of these cells (D). The over-migration phenotype and the increase of glial density caused by the over-expression of *UAS-tkv^QD^* and *UAS-ihog* with *repo-Gal4* was suppressed in the *hep^r75^* mutant discs. Nonetheless, the glial cell division was not significantly altered (E and G). (F) The graph represents the glial density of discs shown in A-E. (G) Histogram showing the % of glial cells in mitosis (Ph3 positive) of discs indicated in A-E. (H)The graph represents the % of discs with glial overmigration show in A-E.

These results further support a model in which synergistic interaction between *hh* and *dpp* signalling stimulate proliferation and glia motility.

### Glial motility stimulation by *dpp* and *hh* signalling depends on JNK function

The results described so far indicate that JNK signalling facilitates glial motility during normal development, as well as in response to damage. Similarly, our observations suggest that *dpp* and *hh* signalling stimulate both the proliferation and the motility of the subretinal glia. We investigate whether JNK mediates some of these functions promoted by *dpp* and *hh* signalling. If the increased motility of glial cells produced by *dpp* and *hh* signalling is mediated by JNK, we expect that the hyperactivity of these signalling pathways might up-regulate JNK signalling in glia cells. To examine this idea, we analysed the activity of *puc2B-Lacz* in discs over-expressing *UAS-dpp* and or/*UAS-hh* in the retinal region under the control of *tub-Gal 80^ts^ GMR-Gal4*. The ectopic expression of *UAS-hh* did not significantly modify the activity of this reporter, while the over-expression of *UAS-dpp* increased both the percentage of glial cells that express the reporter, as well as the proportion of glial that express the reporter at high levels (Figure S13). The co-expression of both factors strengthens the latter effect, though the percentage of glial cell expressing *puc2B-Lacz* is comparable to that observed when *UAS-dpp* was expressed alone (Figure S13). Surprisingly, the autonomous activation of *dpp* (*UAS-tkv^QD^) or hh* signalling *(UAS-ihog)* individually in glial cells with *repo-Gal4* did not significantly affect the activity of JNK-signalling, as assayed with TRE-GFP (Figure S14). Nonetheless, the co-activation of *UAS-tKv^QD^* and *UAS-ihog* under the control of *repo-Gal4* using the same experimental condition drastically elevated the activity of *TRE-GFP* in glial cells (Figure S14).

All together our results suggest that the ectopic expression of *dpp* and *hh* in the eye discs can promote non-autonomous activation of these pathways in glial cells, that in turn stimulates JNK signalling in glia cells. To define whether JNK signalling mediates some of the effects induced by these two signalling pathways, we examined glial proliferation and density in *hep^r75^* mutant discs over-expressing *UAS-tkv^QD^* and *UAS-ihog* under the control of *repo-Gal4*. In this mutant background there is a drastic reduction in the density of glial cells compared to *UAS-tkv^QD^ UAS-ihog repo-Gal4* discs (Figure 9). However, glial cell proliferation was not significantly altered. We also found that the over-migration phenotype caused by the over-expression of *UAS-tkv^QD^ UAS-ihog* with *repo-Gal4* was totally suppressed in the *hep^r75^* mutant background (Figure 9). Therefore, these results are consistent with a model in which the function of JNK is necessary for facilitating the over-migration of glial cells induced when *dpp* and *hh* signalling are over-expressed.

### Glial activity in response to damage is mediated by JNK signalling

We found that wrapping glia, and some perineural glia cells, respond to damage in the retina region by changing in their morphology and behaviour. Next, we investigated whether the JNK signalling pathway, which was highly activated in some of these cells in response to damage, influences some of these effects. To this end, we examine WG morphology and behaviour upon damage induction in different mutant conditions for genes members of the JNK signalling pathway.

To knockdown JNK function specifically in wrapping glial cells, we over-expressed *UAS-puc* or a dominant negative form of Basket under the regulation of *Mz97-Gal4.* In undamaged *Mz97-Gal4 UAS-GFP UAS-puc* discs the density, proportion (WG/Total glial cells), morphology and localization of WG were not altered when compared with control discs (*Mz97-Gal4 UAS-GFP*). However in discs over-expressing *bsk^DN^* under the control of *Mz97-Gal4* or in *hep^r75^* mutant discs, we observed a weak, but statistically significant reduction of the density of WG (Figure 10), although the proportion of these cells was not altered. The down-regulation of JNK signalling in the WG of injured discs, either via the over-expressing *UAS-puc* or *UAS-bsk^DN^*, altered neither the density of these cells nor their location in the apical/middle layers of the discs (Figure 10). However, we did detect a reduction in the proportion of WG (Wg/Total Glial cells) in these discs compared to injured control discs (Figure 10 L). Moreover, although we observed neuronal debris inside these WG cells, they hardly ever developed cytoplasmic projections as those presented in activated WG in control discs (Figure 10). This is most clearly seen in a time-lapse image analysis. We observed that the transformed WG in damaged discs where JNK signalling has been blocked barely produced membrane projection toward the damaged region, compared to control damaged discs (compared video 2 with video 5). Another feature of activated wrapping glial cells was the enlargement of their nuclei. Interestingly, this effect was suppressed when JNK was reduced (Figure 10 M).

**Figure 10.**
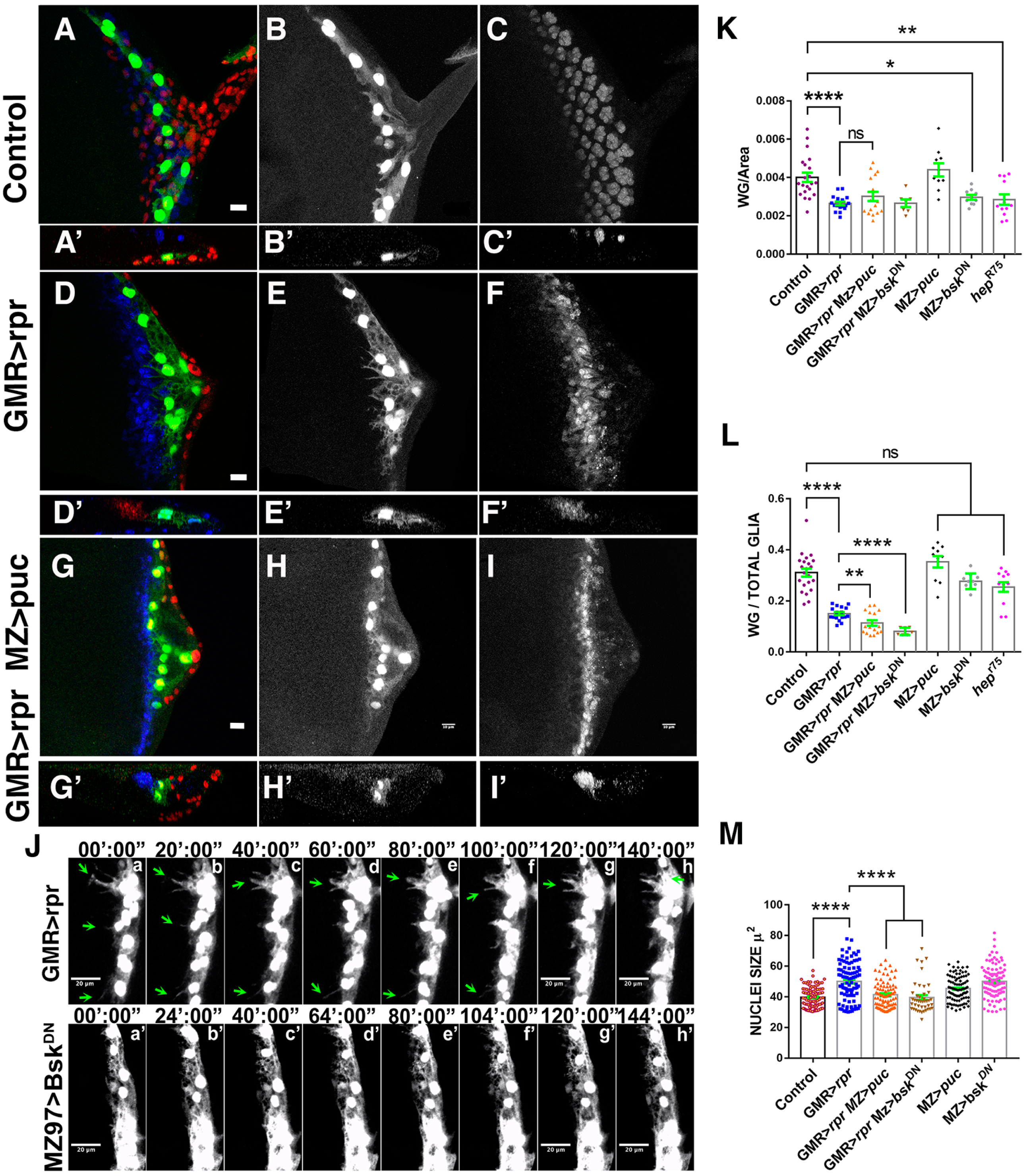
JNK depletion affects the morphology and behaviour of wrapping glial in damaged eye discs. (A-I’) Third instar eye discs stained with anti-Elav (Blue) and anti-Repo (red). (A-I’) *Mz97-Gal4* driving expression of *UAS-GFP* (green) reveals wrapping glial cells in control *UAS-GFP Mz97-Gal4; QUAS-rpr* (A-C’), and damaged eye discs with JNK signalling (*GMR-QF; UAS-GFP Mz97-Gal4; QUAS-rpr,* D-F’) and without this signal (*GMR-QF; UAS-GFP Mz97-Gal4; repo-Gal4 QUAS-rpr/UAS-puc,* G-I’). (A’-C’, D’-F’ and G’-I’) Cross-sections perpendicular to the furrow of eye discs shown in A-I. The apical localization of wrapping glial cells observed in damaged discs is not prevented by the reduction of JNK signalling (compared E’ to H’). However, when the function of JNK signalling is blocked, wrapping glial cells do not produce the large processes found in wrapping glial in damaged disc (compared E to H). (J) Detailed frames from *in vivo* time-lapse analysis (video 5) of eye discs. When JNK is blocked wrapping glial cells do not form celular projections, as it occurs in damaged discs. (K-M) Graphs showing: wrapping (Wg) and perineurial (PN) glial density (K), ratio of wrapping (Wg) and perineurial (PN) glial cells (n° wrapping or perineurial glial cells/n° Total glia) (L), and nuclei size (M).

## DISCUSION

The eye disc contains several glial cells that overall, have equivalent functions to some of the mammalian glia cells found in the PNS (Rodrigues et al., 2011; Klambt, 2009; Yildirim et al., 2019). The unique developmental features of these discs have made them an ideal model in which to establish the molecular underpinnings of glial development and function (Yuva-Aydemir and Klambt, 2011; Silies et al., 2010; Chotard and Salecker, 2007; Yildirim et al., 2019). However, the use of the eye disc as a model system to study glial response to neuronal damage and the mechanisms that might be regulating them, has remained largely unexplored. Glial regenerative response (GRR) is found across many species and may reflect a common underlying genetic mechanism (Hidalgo and Logan, 2017; Kato et al., 2018). Considering that *Drosophila* glia cells have served as an experimental model to gain insights into mammalian glial biology, eye discs might provide an excellent model system to discover evolutionarily conserved signalling networks regulating glia regenerative response.

The eye disc contains different glial cell types including a specialized perineurial glial cell type, the wrapping glia. This glia type is an axon-associated cell that enwraps axons resembling non-myelinating Schwann cells forming Remak fibers in the mammalian PNS (Yildirim et al., 2019). In these animals, nerve injury triggers the conversion of non-myelin (Remak), as well as, myelin Schwann glia to a cell type specialized in promoting repair. During this process these cells undergo large scale changes in gene expression and morphology. In mammals, peripheral nerves owe their regenerative potential to the ability of Schwann glial to convert to cells devoted to repair after injury (Arthur-Farraj et al., 2012; Jessen and Mirsky, 2016). These properties make Schwann cells attractive candidates for regenerative therapies of the injured spinal cord (Xu et al., 1995a; Xu et al., 1995b).

In this work, we have shown that when the neuronal region of the eye disc epithelium is damaged it induces a glial response that consists of an increase in glial migration and division. In addition, we observed that wrapping glial cells, and some perineural glia, undergo morphological changes that confer to them phagocytic activity. This behaviour resembles the regenerative response of non-myelinating Schwann cells (Arthur-Farraj et al., 2012; Jessen and Mirsky, 2016; Jessen et al., 2015). Moreover, we found that upon damage, wrapping glia cells in the eye disc activate JNK signalling.

Similarly, it has been shown that damage in peripheral nerves of mammals induces c-Jun activation in Schwann cells of the PNS (Arthur-Farraj et al., 2012). Therefore, the eye disc is a good model to reveal regulatory mechanisms involved in glial response in the PNS, which might be of relevance to mammalian organisms.

### Activation of JNK signalling in glia cells is necessary but not sufficient for inducing Glial response

In *Drosophila*, the function of the JNK signalling pathway is required for regenerative processes in both the CNS and PNS. In adult and larva brains, the function of Eiger/TNF is involved in triggering glial proliferation upon injury (Kato et al., 2009). In the PNS, the JNK signalling pathway is activated in response to axon injury in damaged neurons, where it regulates injury-induced transcriptional changes (Xiong et al., 2010). The activation of this signalling pathway in the damaged neuron can promote axonal re-growth (Brace and DiAntonio, 2017; Song et al., 2012). In addition, it has been shown that axon self-destruction leads to the activation of Draper/Ced-1/MEGF-10. This protein is an engulfment receptor that promotes clearance of cellular debris in different organisms, including *Drosophila* (Macdonald et al., 2013). In glial cells, Draper binds to TRAF4 and Shark. This promotes cytoskeletal rearrangements (a process essential for phagocytic activity) and at the same time, induces the activation of JNK signalling, leading to the induction of dAP-1-mediated gene transcription, including Draper. The activation of this factor in glial cells facilitates glial engulfment of axonal debris (Lu et al., 2017).

The involvement of JNK in promoting other aspects of glial regenerative response, such as glial proliferation and motility in the PNS remained largely un-explored. The data presented in this work suggest that the function of JNK signalling is required in both the damaged region, as well as in subretinal glial cells to promote glial motility but not proliferation. This effect seems to be only permissive, as the ectopic activation of JNK signalling was not sufficient for inducing glial motility. Our results also indicate that the activation of JNK in subretina glial cells in response to damage, was not mediated by the ligand Eiger. An alternative mechanism to explain how JNK signalling might be activated in the damaged eye is via Drapper (Macdonald et al., 2013). We found; however, that conversely to what it occurs in other tissues such as in antennal segments, in damaged eye discs, Drapper was not ectopically expressed (data not shown) and likely not required for inducing JNK in damaged eye discs, although further experiments are required to confirm this.

Damage in peripheral nerves of mammals provokes the activation of the transcription factor c-Jun (Arthur-Farraj et al., 2012). This early event in the regenerative response plays a fundamental role in regulating the major aspects of injury response. It determines the formation of regeneration tracks and myelin clearance and is required for efficient Schwann cell de-differentiation. This later process activates a repair program and transforms these cells into cells specialized to support regeneration (Arthur-Farraj et al., 2012). The repair Schwann cell is a transient cell state required for repairing injured tissue (Jessen and Mirsky, 2016). Similarly, our results suggest that in the eye disc, damage induces the activation of JNK signalling in wrapping glial and subsequently, these cells undergo various morphological and behavioural changes. We show that the reduction of JNK signalling in transformed WG partially suppresses some of the effects induced by damage, for instance the formation of the long membranes projection observed in transformed WG was inhibited. However, our data indicate that the ectopic activation of JNK signalling in WG in control undamaged discs was not sufficient for inducing the WG transformation, suggesting that other signal pathways must be involved in regulating this process. Altogether, these results suggest that the combination of different signals is necessary to promote the transformation of WG cells in response to damage.

### *dpp* and *hh* signalling promotes JNK activation for facilitating glial motility

During the development of the eye, *dpp* and *hh* pathways drive a wave of neuronal differentiation across the eye disc. In this process, the function of these signals is partially redundant, indicated by the fact that neuronal differentiation was only blocked when both pathways were simultaneously eliminated (Curtiss and Mlodzik, 2000). The coordination of glial migration and growth of the eye imaginal disc is also mediated by *dpp* and *hh* signals (Rangarajan et al., 1999). The cells of the eye disc secrete Dpp and Hh that act on perineurial glial cells, promoting their migratory abilities and inducing their proliferation. Dpp promotes both the proliferation and motility of the glia, whereas Hh appears to promote only their motility (Rangarajan et al., 1999). Both pathways collaborate during this process, since the simultaneous elimination of both pathways have stronger effects that when each signal was blocked individually (Rangarajan et al., 1999). Our data indicate that as it occurs during normal development, both pathways collaborate for controlling glial behaviour in response to damage.

The down-regulation of *dpp* in the injured region was not sufficient for modifying glial response. Only the simultaneous silencing of *dpp* and *hh* in the damage region altered glial behaviour in our experimental condition. As we previously mentioned, this might be due to the weakness of the loss of function effects caused by RNAis. Alternatively, both signalling pathways might have a redundant function in promoting glial response; as previously shown during neuronal differentiation in eye discs (Curtiss and Mlodzik, 2000). However, we have observed that when the function of *dpp* was eliminated in glial cells of injured discs, glial response was strongly supressed. Whereas the diminution of *hh* function in glial cells did not significantly affect this response. These results argue against the idea that these pathways have a redundant function regulating this process. Nevertheless, since both signalling events collaborate to promote glial response, and the effects caused by the simultaneous elimination of both signalling pathways were stronger than when each pathway was individually altered, we cannot rule out that they might have a partially redundant function inducing glial response.

We have shown that part of the function of *dpp* and *hh* signalling inducing glial response is mediated by JNK signalling. The over-expression of *hh* and *dpp* leads to high activity of JNK signalling in glial cells. Previous work has shown that in wing discs, that localized JNK activity is established by Hedgehog signalling through the induction of TNF receptor-associated factors (dTRAF1) expression. dTRAF1 is known to act upstream in activation of the JNK signalling pathway through association with the MAP kinase kinase kinase kinase (Liu et al., 1999; Willsey et al., 2016). Although further experiments are required to elucidate the mechanisms by which *dpp* and *hh* activate JNK signalling in glial cells, it is possible that in the damaged eye, Hh and Dpp together facilitate the up-regulation of dTRAF that, in turn, would activate JNK in glial cells.

Our work reveals that the eye disc is a powerful model system in which to define the mechanisms that control glical cell migration, proliferation and activity transformations which occur in response to neuronal damage. The results presented in this work lead us to propose a mechanism describing how the long-range signalling pathways of *dpp* and *hh* might promote the glial response to neuronal damage. We have found that in the damaged cells of the retina, *dpp* and *hh* are ectopically expressed and in turn, this non-autonomously activates both signalling pathways in glial cells to promote their proliferation and induce the activity of JNK pathway to facilitate their motility (Figure 11).

**Figure 11.**
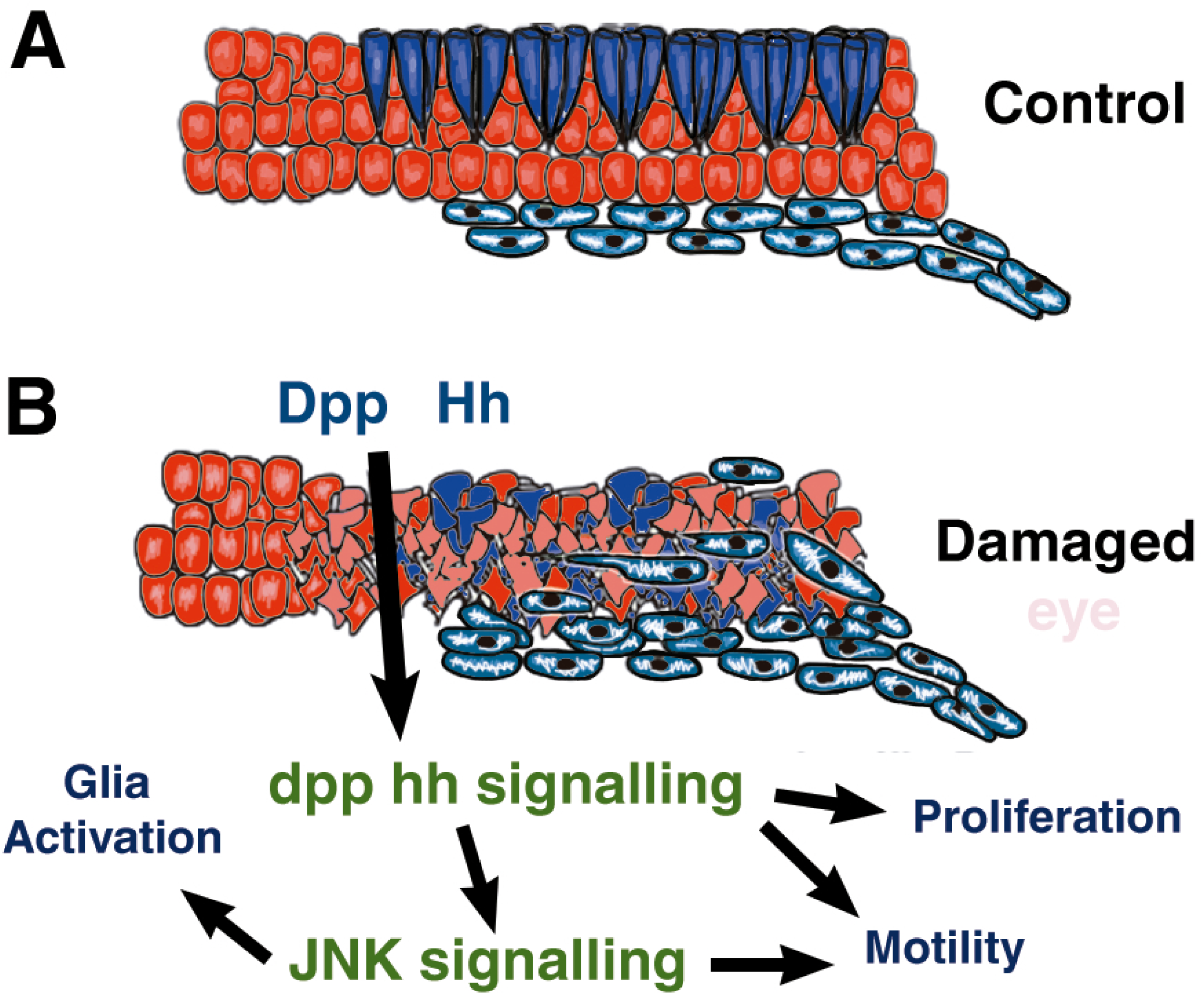
Model. Model representing the signalling network involved in the activation of glial response in damaged eye discs

## Materials and Methods

### *Drosophila* stocks and genetics

The following stocks were used:

*UAS* lines: *UAS-GFP (II and III), UAS-dpp (III), UAS-reaper (X),* (described in Flybase, Bloomington Drosophila Stock Center) *UAS-dpp^RNAi33^* (Bloomington Drosophila Stock Center: 33618)*, UAS-dpp^RNAi2^* (Kyoto DGGR, 9885R-3)*, UAS-dpp^RNAiInt^ (UAS-dppshmiR)* (Haley et al., 2008)*, UAS-hep^CA^ (II) (III)* (Adachi-Yamada et al., 1999), *UAS-tkv^QD^* (Nellen et al., 1996), *UAS-tkv^DN^* (Haerry et al., 1998), *UAS-brk* (Jazwinska et al., 1999), *UAS-puc2A* (Martin-Blanco et al., 1998), *UAS-bsk^DN^* (Adachi-Yamada et al., 1999), *UAS-hh-GFP* (II)(Torroja et al., 2004), *UAS-ihog* (VDRC stock: 29897), *UAS-hh^RNAi^* (VDRC stock:17874), *UAS-boi^RNAi^* (VDRC stock:3060), *UAS-ihog^RNAi^* (VDRC stock: 29897) *UAS-rbf^280^* (Xin et al., 2002) and *UAS-microRNARHG (II)* (Siegrist et al., 2010).

Mutants: *eiger^1^/Cyo, eiger^3^/Cyo*, and *hep^r75^/FM7*. All of these stocks have been previously described in FlyBase.

The reporter lines: *puc-LacZ* line (Martin-Blanco et al., 1998), *dpp-LacZ* and *hh-LacZ* (described in Flybase) and TRE-DsGFP (Chatterjee and Bohmann, 2012). We used the Gal4 lines: *GMR-Gal4 tub-Gal 80^TS^ /CyO, tub-Gal80^TS^; repo-Gal 4/TM6B, c527-Gal4* and *Mz97-Gal4* (Hummel et al., 2002).

We used the *QF/QUAS* lines: *GMR-QF; tub-Gal80^ts^/Cyo; repo-Gal4/TM6B, QUAS-rpr repo-Gal4/TM6b* and *QUAS-rpr /TM6b* (*Potter and Luo, 2011; Potter et al., 2010*).

Stocks and crosses were maintained on yeast food at 25°C or 17°C before and after the transitory inhibition of Gal80^TS^ at 29°C.

### Genetic analysis

This analysis was performed by crossing *UAS-rpr*: *GMR-Gal4 tub-Gal80^ts^ /CyO*, and *GMR-Gal4 tub-Gal80^ts^ /CyO* to the following stocks:

UAS-GFP
*UAS-bsk^DN^*
*UAS-puc2B*
*UAS-dpp^RNAi2^,*
*UAS-dpp^RNAi2^; UAS-hh^RNAi^*
*_UAS-dpp_^RNAiInt^*
*UAS-dpp^RNAi33^*
*UAS-hh^RNAi^*
*UAS-dpp*
*UAS-hh-GFP*
*UAS-hh;UAS-dpp*
*puc-LAcZ*
*dpp-LacZ*
*hh-LacZ*

We also crossed *UAS-rpr*: *If /CyO; UAS-hh^RNAi^/TM6B* to:

GMR-Gal4 tub-Gal80^ts^ /CyO; UAS-dpp^RNAiInt^/TM6B
GMR-Gal4 tub-Gal80^ts^ /CyO; UAS-dpp^RNAi33^/TM6B

To study the effect of ectopic expression of the different *UAS* lines used in our analysis *en*–*Gal4 tub-Gal80^TS^ UAS-X* larvae (where X indicates the different transgenes used in our assay) were raised at 25°C until 132±12 hours AEL, at which point the larvae were shifted to 29°C for 24 hours and then shifted back to 25°C. We analysed the effects caused by the overexpression of the different UAS lines immediately after the end of the shift to 29°C (T0), and 24 hourss later (T1) (Figure S1).

### Immunocytochemistry

Immunostaining of the wing discs was performed according to standard protocols. The following primary antibodies were used: rabbit anti-Phospho-histone 3 1:200 (Cell Signaling Technology), rabbit anti-cleaved Dcp1 1:200 (Cell Signaling Technology), mouse anti-Patch 1:500 (Gift from Isabel Guerrero), mouse anti-ß Galactosidase 1:200 (Promega), anti-ß Galactosidase 1:500 (Cappel), rat anti-Phospho-Mad 1:100 (from Ginés Morata). Mouse anti-Repo, rat anti-Elav 1:200, were obtained from the Developmental Studies Hybridoma Bank at the University of Iowa. Secondary antibodies (Molecular Probes) were used at dilutions of 1:200.

Imaginal discs were mounted in Vectashield mounting fluorescent medium (Vector Laboratories, Inc.).

### EDU staining

Eye imaginal discs were dissected in Phosphate Buffer Saline (PBS) and then incubated in a solution with 5-Ethynyl-20-deoxyuridine (EdU) for 1 hour at room temperature to label cells in S phase. Alexa Fluor detection was performed according to Click-iT EdU Fluor Imaging Kit (Invitrogen)

### Puc reporters construction

We selected two different regions of the regulatory region of the gene *puckered* puc-1 and puc-2, see Figure S7) on the basis of the published open chromatin profile of eye discs during tumour development. The selected region were amplified using KOD enzyme (Novagen) by PCR, using the following primers:

PUC1: Forward: CAGTAAGCTTGCCGTCAACTTTTATCTGCCAACG
Reverse: CAGTAGATCTCGGGCTAATTGGACTGGGGTTCAA
PUC2: Forward: CAGTAAGCTTGGGGTGGCAATGACTCACAATAGG
Reverse: CAGTAGATCTCTGCAAAGATACATGCGGATCGG

The PCR products were cloned into the attB-hs44-nuc-lacZ vector, using the HindIII and BglII restriction endonucleases (NEBiolabs).

The two sequences (*puc1* and *puc2*) were further subdivided into smaller overlapping fragments. The *puc1* region of 2782 bp was divided into three fragments: *puc1A* (978 bp), *puc1B* (938 bp) and *puc1C* (839 bp); while the *puc2* region of 1636 bp was sub-divided into two fragments: *puc2A* (897 bp) and *puc2B* (578 bp). To this end, PCR were performed using the aforementioned attB-hs44-puc1-nuc-lacZ and attB-hs44-puc2nuc-lacZ constructs as template. We added targets for the restriction enzymes BglII and HindIII to all the fragments generated, and then cloned into the attB-hs44-nuc-lacZ vector using these restriction sites

### Quantitative analysis

Images were processed using the ImageJ software (NUH,Bethesda,USA).

We calculate the glia density as the ratio between the number of glial cells, as detected by the expression of Repo, and the size of the region posterior to the morphogenetic furrow in µm^2^ (Repo positive cells/size of the area in µm^2^). Eye discs were measure using ImageJ.

Glial cell division was calculated as the ratio between the number of PH3-positive glial cells and the size of the region posterior to the morphogenetic furrow in µm^2^ (PH3-positive glial cells /size of the area in µm^2^), or as the percentage of PH3-positive glial cells (PH3-positive glial cells /Total glial cells *100).

The percentage of glial cells incorporating EdU was calculated dividing the number of glial cells incorporating EdU by the total number of glial cells.

To define the cell cycle phases of glial cells we used the Drosophila FUCCI system (Fly-FUCCI). Cells in G1 phase are marks in Green, cells in S-phase in red, and cells in G2 phase in yellow. Fly-FUCC was expressed under the control of *repo-Gal4*, and the percentage of glial cells in the different phases of cell cycle was calculated dividing the glia cells labelled in the different phases of the cell cycle by the total number of glial cells in each eye discs analysed.

We calculate the density of wrapping glial cells, dividing the number of glial cells expressing *UAS-GFP* under the control of *Mz97-Gal4* by the area of the region posterior to the morphogenetic furrow in µm^2^.

The proportion of wrapping glia cells in each eye discs analysed was calculated as the ratio between the number of glial cells expressing *UAS-GFP* under the control of *Mz97-Gal4* and the total number of glial cells (*Mz97-Gal4 UAS-GFP* glial cells */*Repo positive cells glial).

Nuclei size was automatically calculated using the ImageJ software (NUH,Bethesda,USA). We adjusted threshold intensity to visualize the nuclei of wrapping glial cells expressing *UAS-GFP* under control of *Mz97-Gal4*. Then, we automatically calculated the area size of each nuclei using the Area option in Set Measurements. We exclude the areas corresponding to the fusion of two or more glial cells.

The percentage of Glial cells expressing the reporter puc2B was calculated dividing the number of glial cells that expressed puc2B-LacZ at detectable levels by the total number of glial cells in each eye.

To calculate the number of glial cells that expressed *puc2-Lacz* at high levels, we used the threshold tool of imageJ to interactively set lower and upper threshold values (155-250). We considered as high –expressing *puc2-LacZ* glial cells those cells that expressed the reporter above the lower level (155).

To calculate the motility of glia cells we selected several points of the glial front at the start of the movie and the distance to the closest perpendicular point at the end of the movie was measured. If the end point is closer to the MF than the start point, then the distance covered is considered positive. If opposite, the distance is considered negative (p value = 0.0172 n = at least 3 independent movies).

### In vitro culture

Imaginal discs were cultured as described (Aldaz et al., 2010)

### Statistical analysis

For statistical test applied to each experiment, n and p values, please see supplementary Tables. p-values shown on the graphs are indicated with the following asterisk code:*p<0.5; **=p<0.01;***=p<0.001;****=p<0.0001.

The error bars indicate the Standard Error of mean (SEM).

### Microscopy

Images were captured using a Confocal LSM510 Vertical Zeiss and processed with ImageJ or Adobe Photoshop CS4.

## Acknowledgements

We thank Jose Felix de Celis, and Luis Alberto Baena for providing reagents and useful discussion. Claire Hills for helping to improve the manuscript. We are very grateful to Gines Morata, Isabel Guerrero, Hermann Steller, Sergio Casas, the Bloomington Stock Center and the Developmental Studies Hybridoma Bank for providing fly strains and antibodies. This work was supported by a grant obtained from Fundación Ramón Areces. Sergio B. Velarde was supported by The National Council of Science and Technology from México (CONACyT) and by a fellowship from the Ignacio Larramendi fundation and Álvaro Quevedo from “Ayudas para la contratacion de Investigadores Predoctorales”, Consejeria de Ciencia. Universidades e innovacion, Comunidad Autonoma de Madrid.

## Supplementary Figures

**Figure S1.**
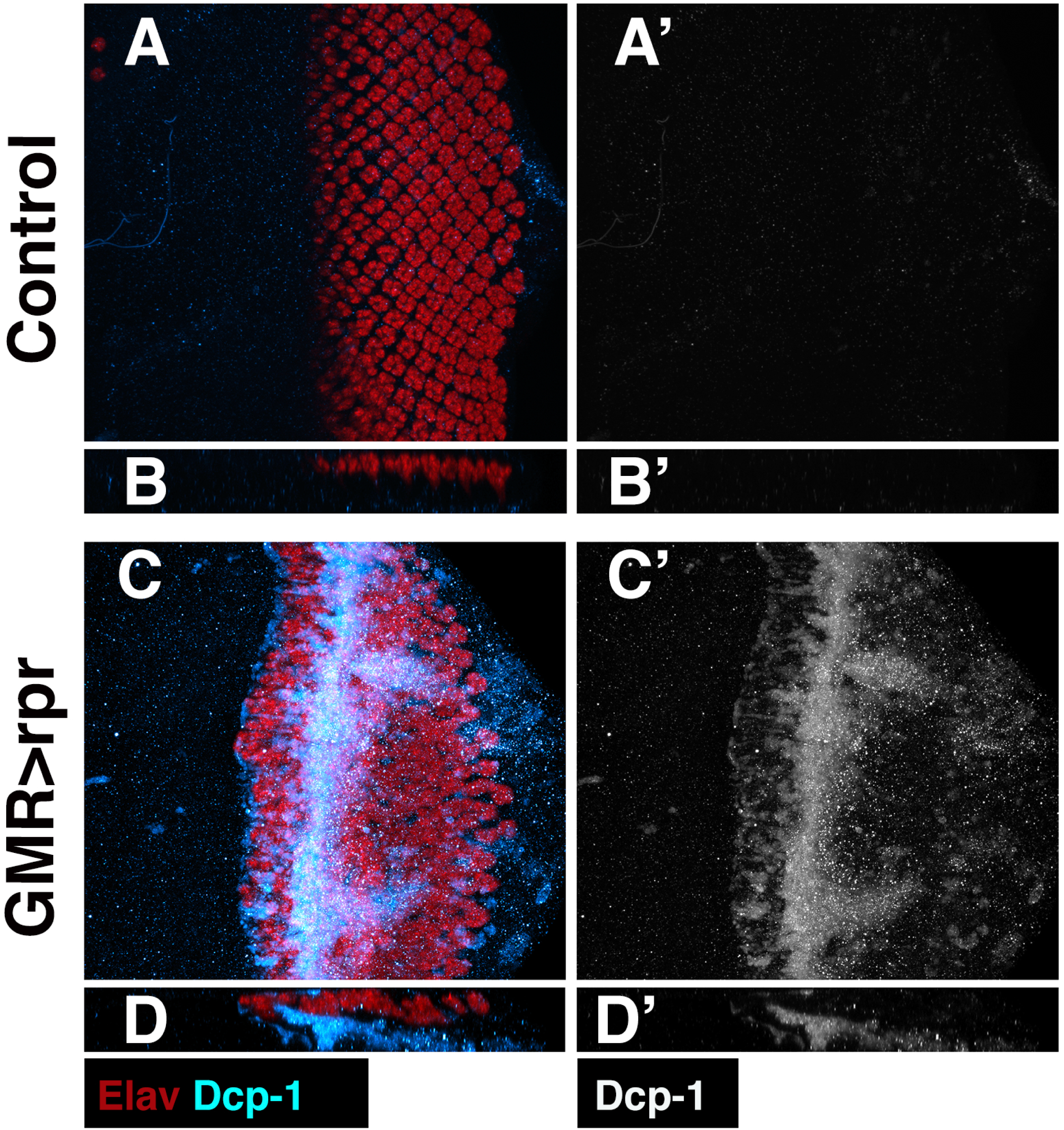
*rpr* expression induces apoptosis in the eye imaginal disc. (A-B’) Third instar eye discs stained with anti-Elav (red) and *Drosophila* Caspase 1 (Dcp-1) (blue in A and B and grey in A’ and B’). (A-A’) Wild-type adult eye, the photoreceptors (red channel) are located at the apical layer of the eye disc. (B-B’) Third instar *GMR>rpr* eye disc. Cell death is induced in the posterior region of the eye disc as assayed by Dcp-1 staining. The X–Z projections below each panel show a cross-section of the epithelium perpendicular to the furrow.

**Figure S2.**
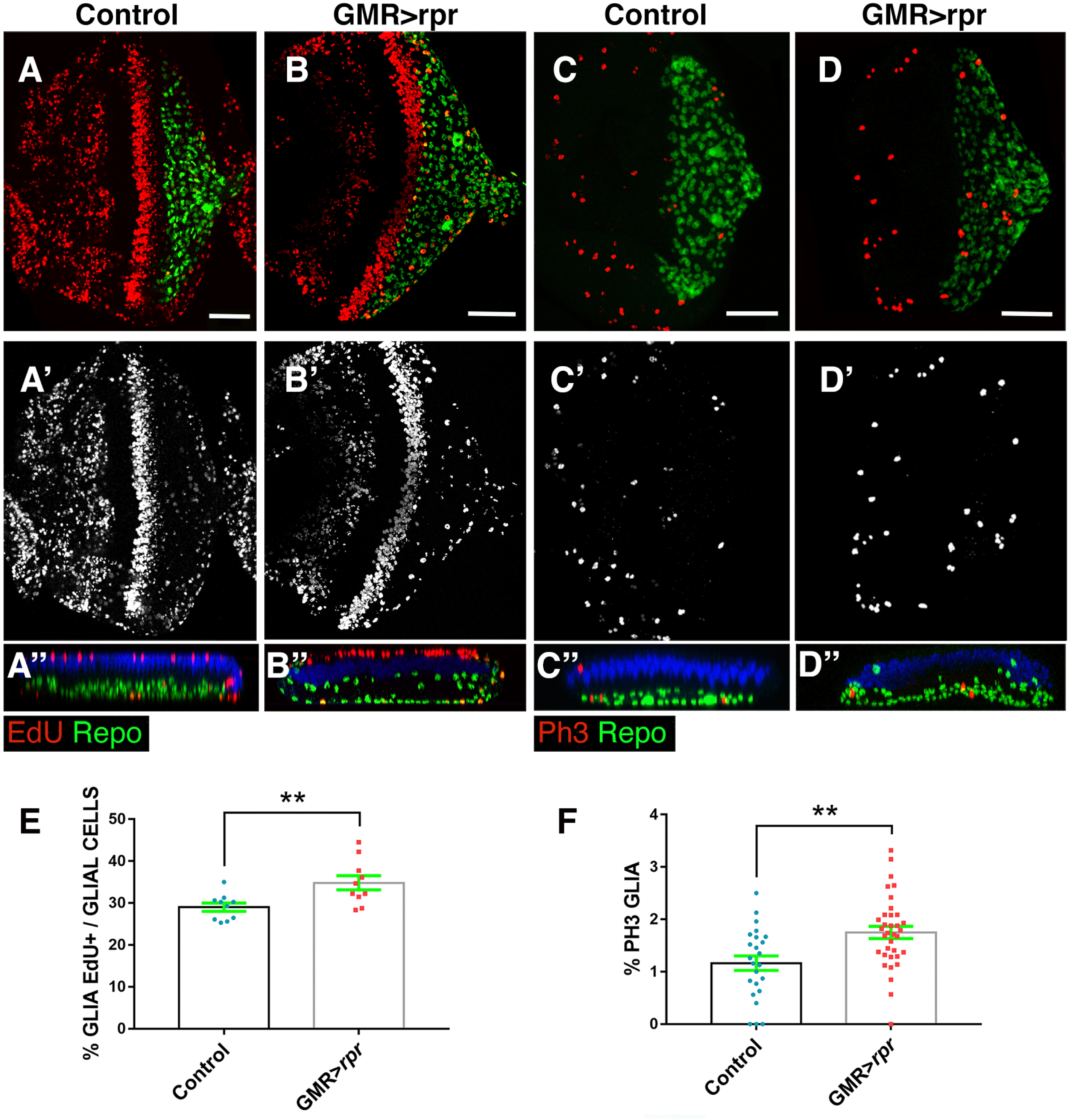
Glia proliferation in damaged discs. (A-F) Third instar control (A and C) and *UAS-rpr/+; GMR-Gal4 tub-Gal80^ts^* eye discs (B and D), labelled with anti-Elav (blue) and anti-Repo (green) (A-D’’). (A’’-D’’) X–Z projections show cross-sections of the eye discs epithelium perpendicular to the furrow of the discs shown in A-D. (E and F) Graphs showing % of glia cells incorporating EDU (E), % of glial in mitosis (PH3-positive Glia /Total glial cells*100) (F). (A-A’’) EDU incorporation in control wt discs (red). (B-B’’) In *UAS-rpr; GMR-Gal4 tub-Gal80^ts^* larvae third instar eye discs raised at 25 °C for 72hr for inducing genetic ablation, we observed significant increase in the percentage of glial cells incorporating EDU (E). (C-C’’) (C) *GMR-Gal4 tub-Gal80^ts^* eye disc stained with antibodies against the mitotic marker eye discs the number of mitotic glial cells increases. Here and in the rest of figures Statistical analysis is shown in supplementary Tables. Scale bar 50μm

**Figure S3.**
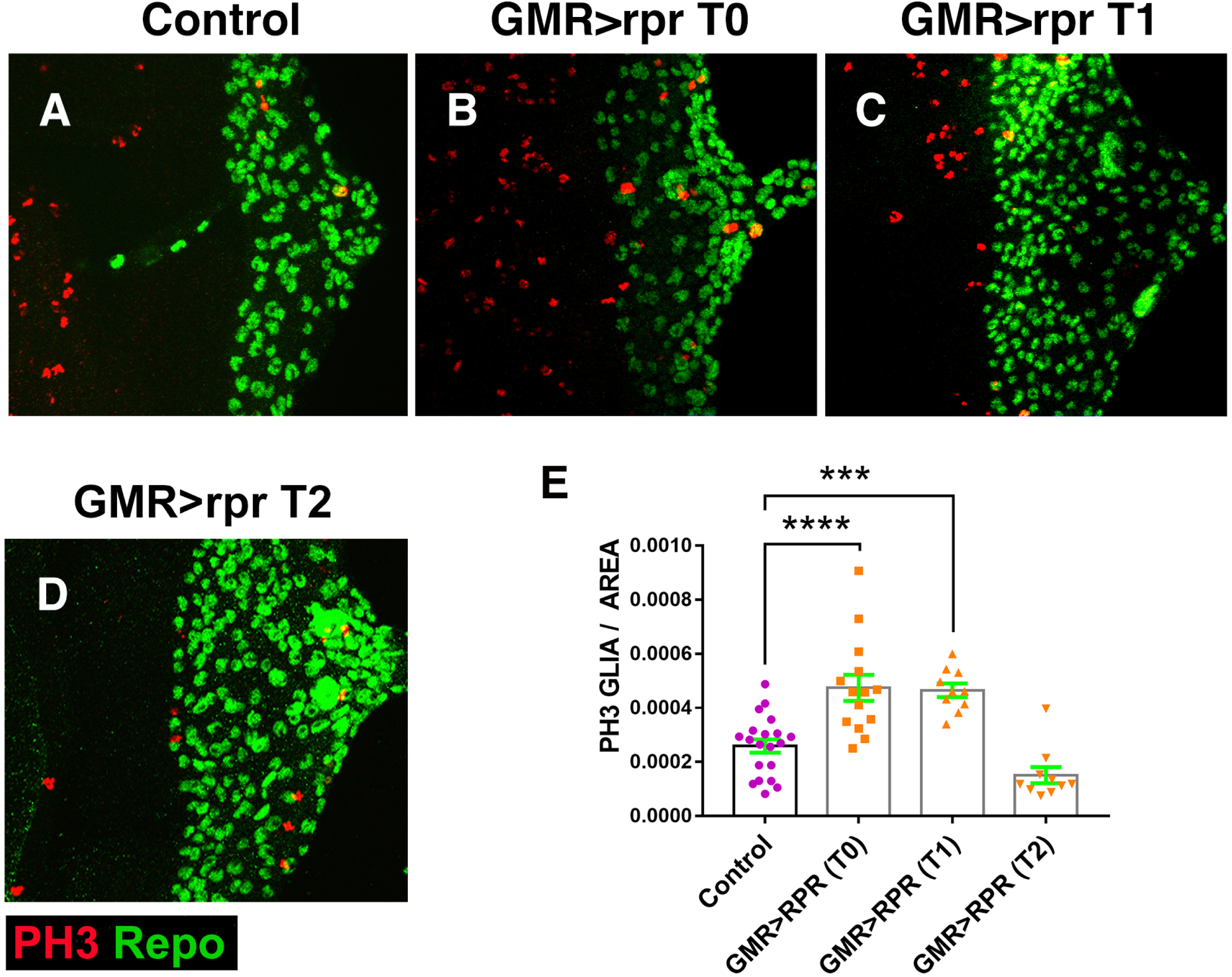
Glial proliferation in response to damage at different times after damaging eye discs. (A-D) Third instar eye discs stained with anti-repo (green) and anti-PH3 (red). Control undamaged disc (A) and *UAS-rpr/+; GMR-Gal4 Tub-Gal80^TS^* eye discs analysed at different times after damage (C-D). We observe a significant increase of mitotic glial cells in damaged *UAS-rpr/+; GMR-Gal4 tub-Gal80^TS^* eye discs immediately after ablation (T0), and after 24 h (T1), but not after 48 h (T2) of recovery. (E) The graph shows the glial mitotic index of control discs compared to discs analysed immediately after damaging (T0), after 24 (T1) and 48 h (T2) of recovery.

**Figure S4.**
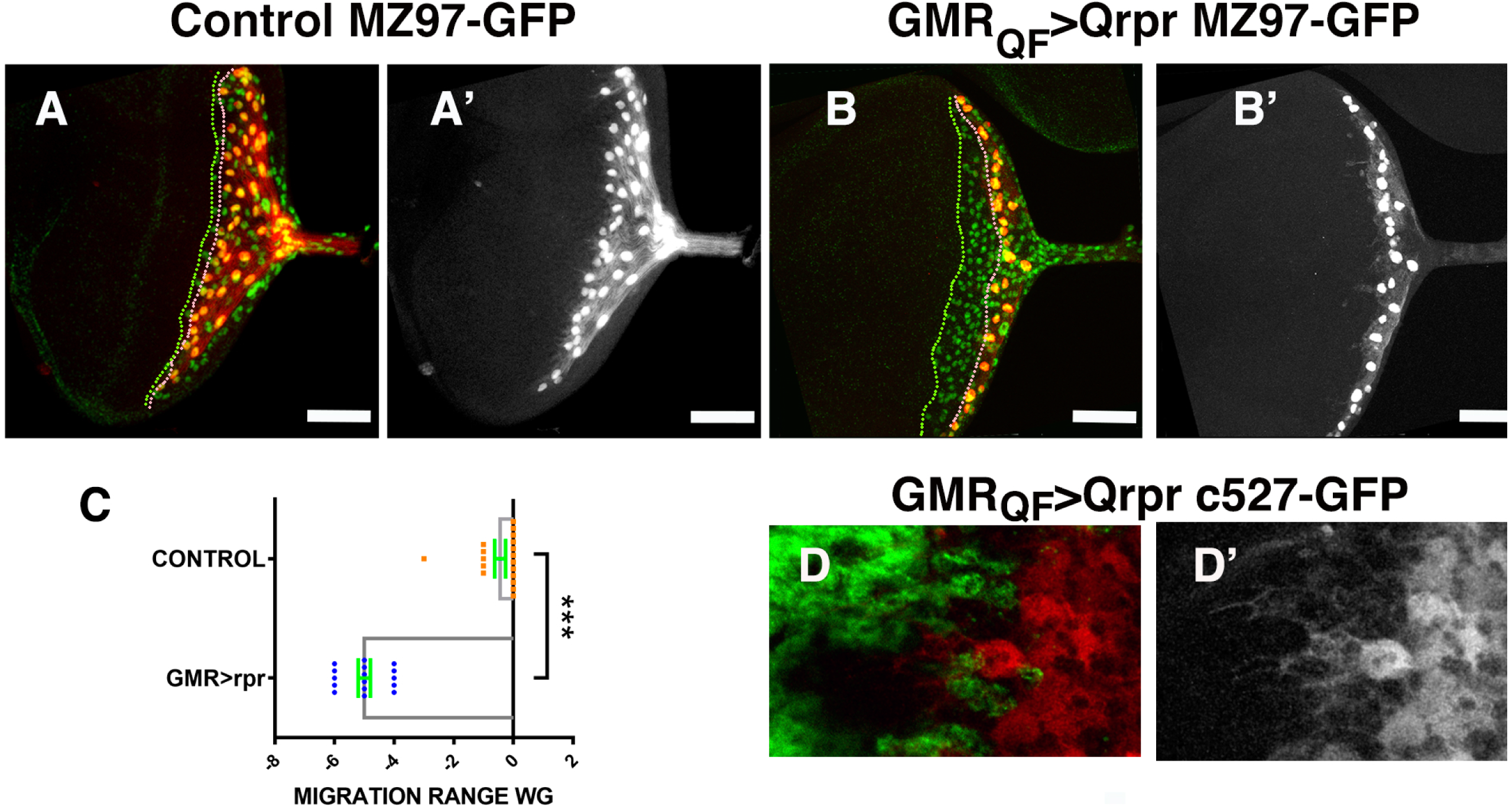
Wrapping and perineurial glial cells generate new projections in direction to the damage area. (A-B’) Third instar eye discs stained with anti-repo (green). Mz97-Gal4 driving expression of *UAS-GFP* reveal wrapping glial cells in control *UAS-GFP Mz97-Gal4; QUAS-rpr* (red in A-A’) and in damaged *GMR-QF; UAS-GFP Mz97-Gal4; QUAS-rpr* (red in B-B’) discs. (A-B’) Projections of the entire confocal stack of control (A-A’) and damaged eye discs (B-B’). Wrapping glial cells in control discs produce large processes that follow the photoreceptor axons toward the brain through the optic stalk, in *GMR-QF; UAS-GFP Mz97-Gal4; QUAS-rpr* damaged we do no observed these projections. (B-B’). Dotted green line shows the anterior border of perineurial glial migration, that in control discs coincides with the anteriormost row of wrapping glial (dotted pink line). In damaged discs wrapping glial cells are located in a region more posterior than the perineurial glial cells. (C) Graph shows the relative position of the anterior border of glial migration with respect to the anteriormost row of wrapping glial. (D-D) *c527-Gal4* drives expression of *UAS-GFP* (red) in perineurial glial cells in damaged *GMR-QF; UAS-GFP c527-Gal4; QUAS-rpr* eye discs. Some of these cells extend large processes towards the damaged region. Elav staining is shown in green.

**Figure S5.**
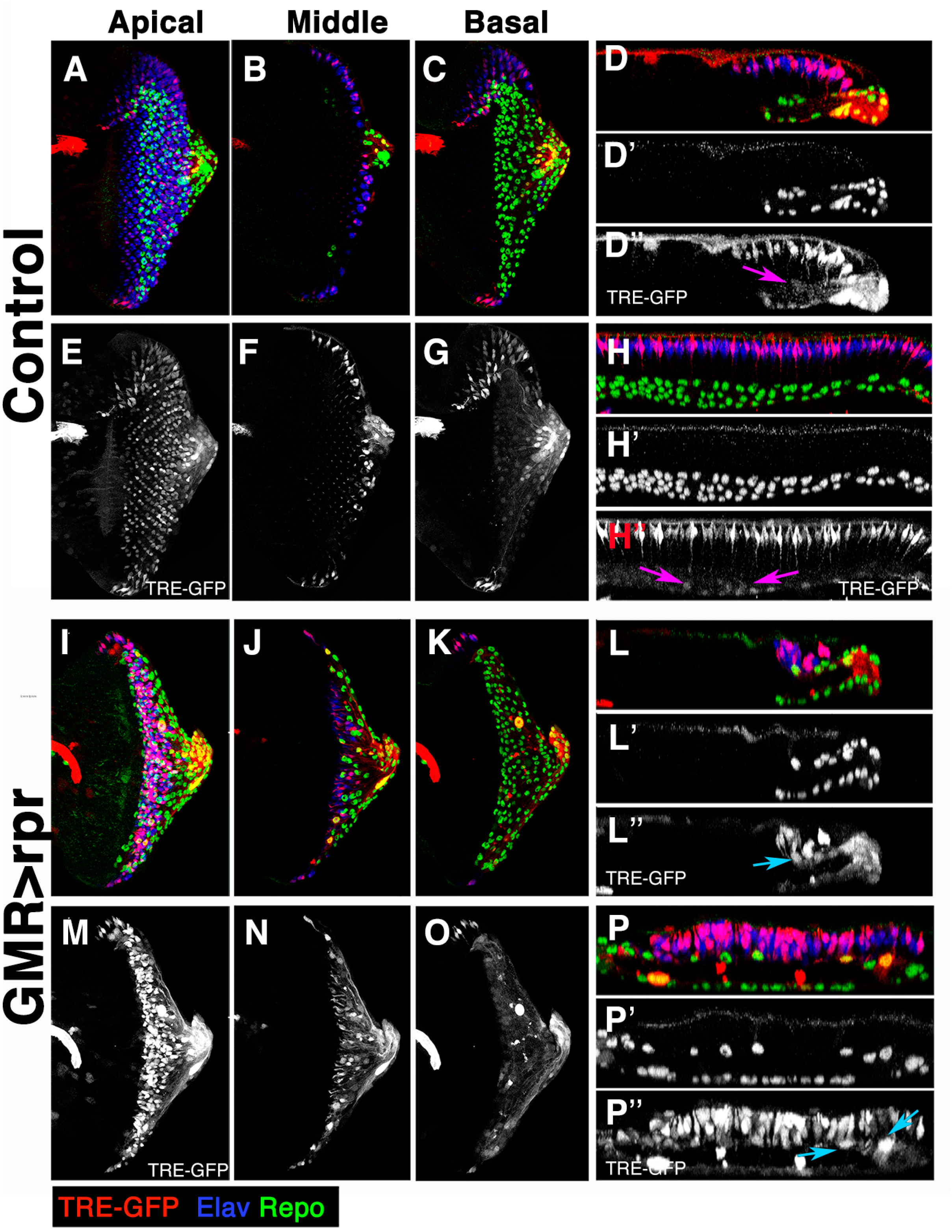
JNK pathway is ectopically activated in glial cells response to damage in the retina. (A-P’’) Third instar eye discs stained with anti-Repo (green and grey in D’, H’, L’ and P’) and anti-Elav (blue). The *TRE-GFP* expression (red and grey in D’’, E-G, H’’, M-O, and P’’) in control *(GMR-Gal4 tub-Gal80^TS^*) (A-H’’) and damaged *UAS-rpr/+; GMR-Gal4 tub-Gal80^TS^ /TRE-GFP* eye discs (I-P’’). Apical (A, E, I and M), Middle (B, F, J and N) and Basal (C, G, K and O) layers. Cross-sections perpendicular (D-D’’ and L-L’’) and parallel (H-H’’ and P-P’’) to the furrow. Pink arrows point glia with a low expression of *TRE-GFP*, while blue arrows point glia and cells with high JNK pathway activation. Glial cells with higher levels of *TRE-GFP* are preferentially located in apical and middle layers of the damaged discs (M, N, L’’ and P’’).

**Figure S6.**
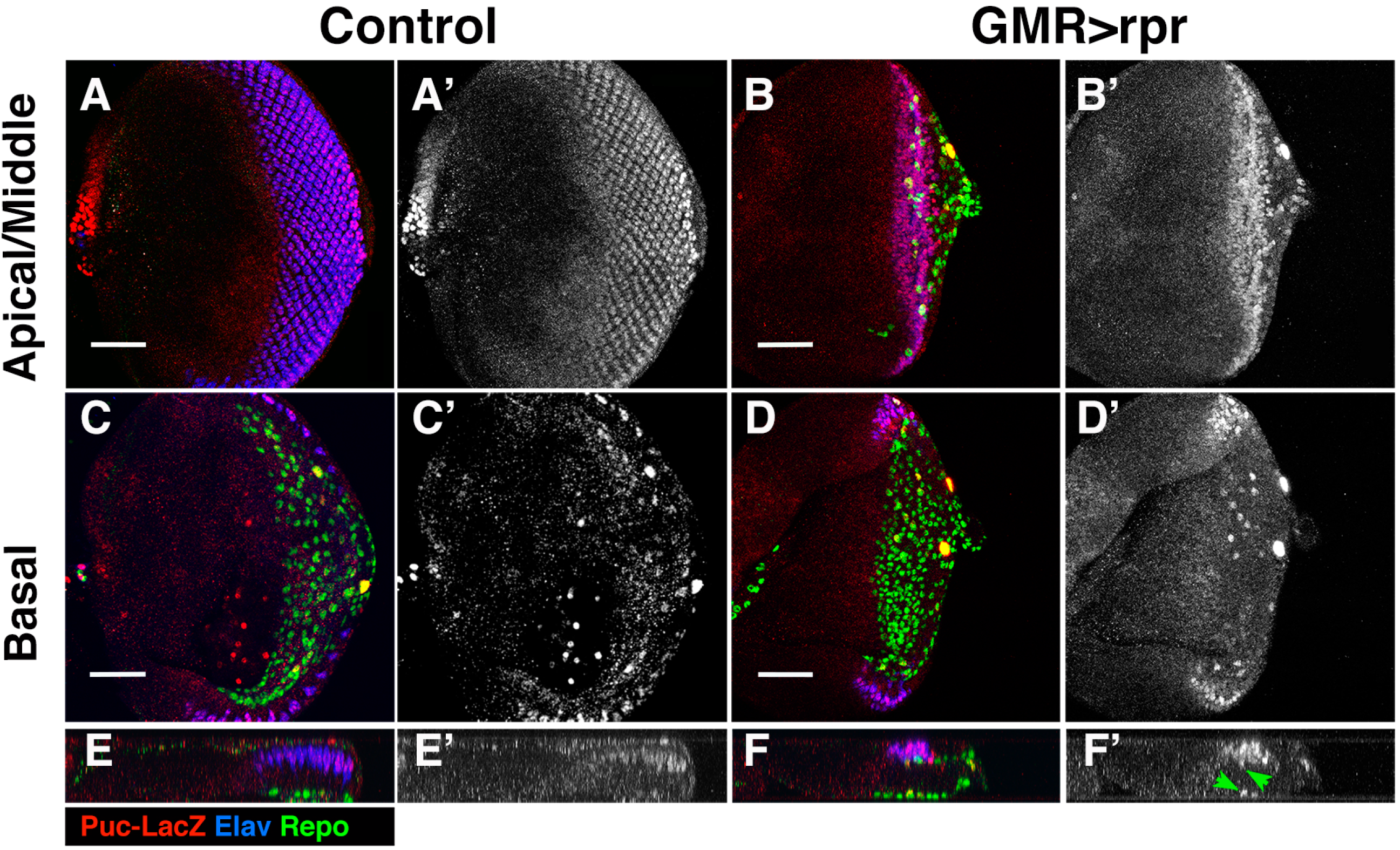
JNK signalling is activated in response to damage. (A-F’) Third instar eye discs stained with anti-Elav (blue), anti-Repo (green) and anti-B-galactosidase (red) to reveal the activity of the *puc-LacZ* reporter, in control *(GMR-Gal4 tub-Gal80^ts^/+; puc-LacZ/+*) (A-A’, C-C’ and E-E’) and damaged *UAS-rpr/+; GMR-Gal4 tub-Gal80^ts^ /+; puc-LacZ/+* discs (B-B’, D-D’ and F-F’). Apical/Middle (A-B’) and basal (C-D’) layers of control (A-A’’ and C-C’’), and damaged (B-B’ and D-D’) eye disc. (E-E’ and F-F’) Y–Z projections of cross-sections perpendicular to the furrow of control (E-E’) and damaged (F-F’) discs.

**Figure S7.**
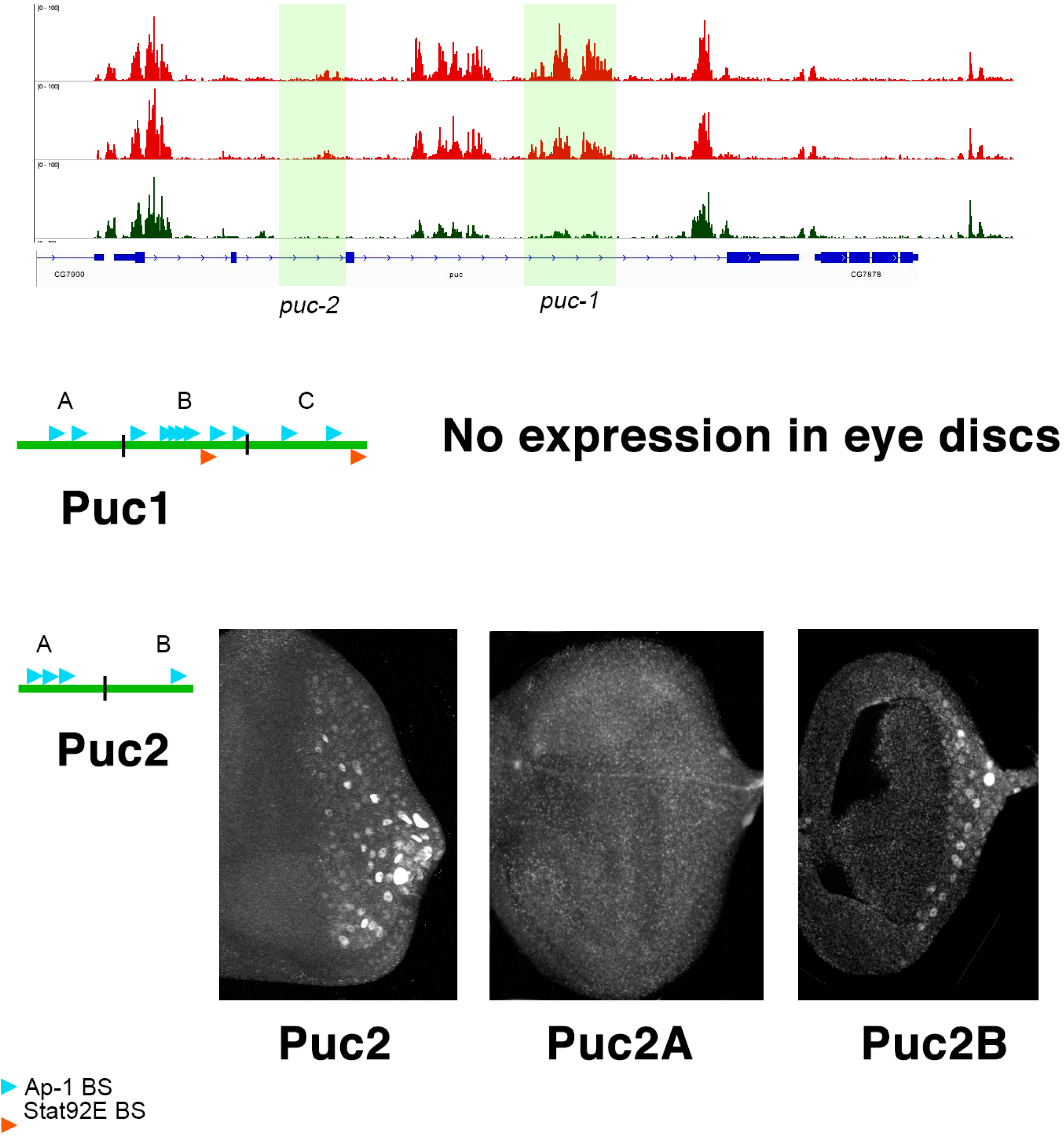
*puc* reporter. Open chromatin profile of the regulatory region of the gene *puc* in the eye discs during tumour development. We selected two different regions (puc-1 and puc-2), that were used to construct *lacZ* reporters. The *puc1* region (2782 bp) was sub-divided into three fragments: *puc1A* (978 bp), *puc1B* (938 bp) and *puc1c* (839 bp); while the puc2 region (1636 bp) was divided into two fragments: *puc2A* (897 bp) and *pucB* (578 bp). The region *puc1* was not sufficient to drive the expression of *lacZ* in the eye discs. *puc2-LacZ* reporter was specifically expressed in glial cells, and a sub-fragment of this element (*puc-2B*) reproduced this pattern of expression.

**Figure S8.**
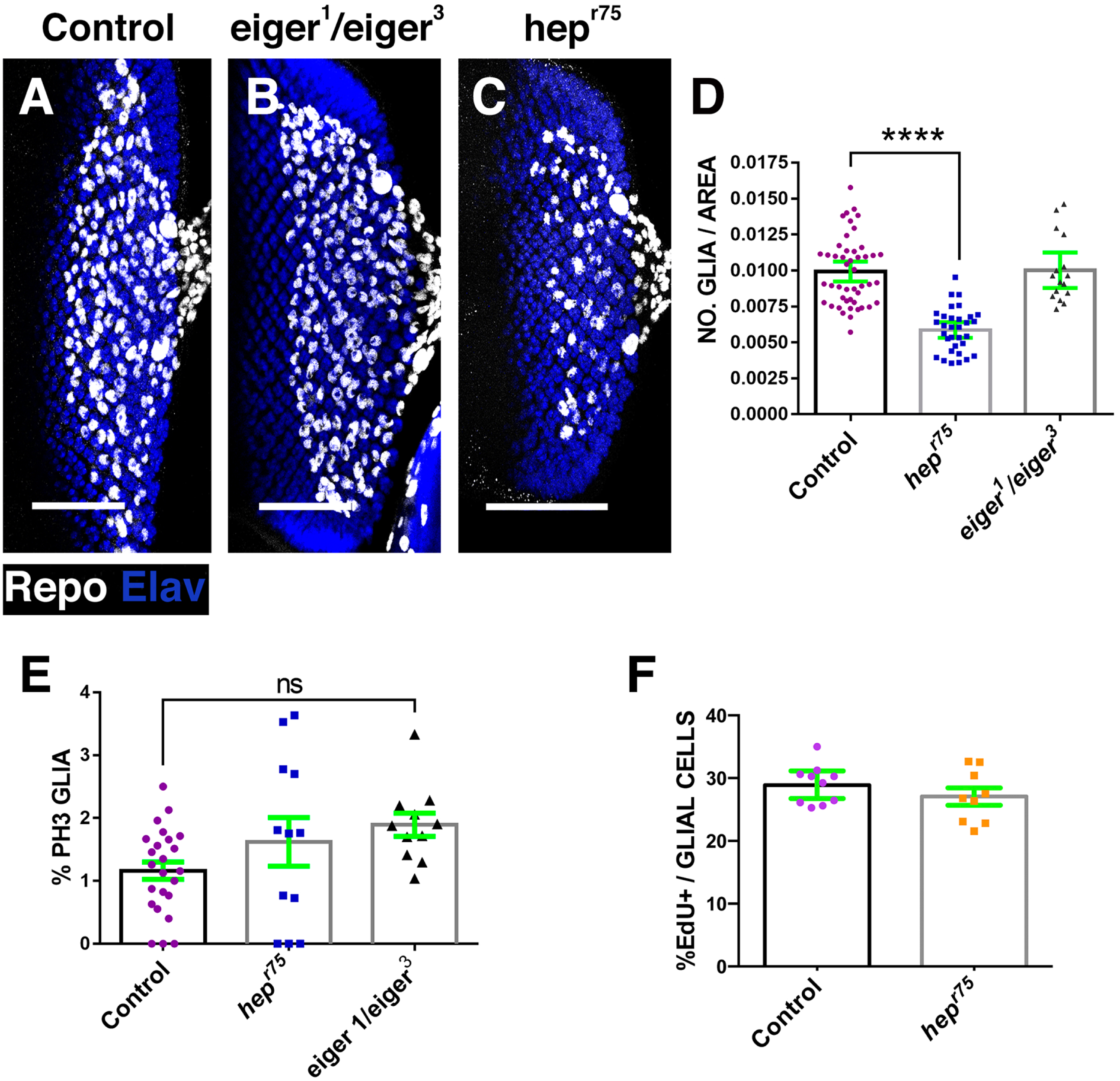
*eiger* is not required during the development of glial cells in the eye disc. (A-C) Projections of confocal images of third instar eye discs stained with anti-Elav (blue) and anti-Repo (white). Control undamaged *GMR-Gal4 tub-Gal80^ts^* (A), *eiger^1^/eiger^3^* (B) and *hep^r75^* mutant eye discs. (D-F) Graphs show glial cell density (n° glial cells/ area) (D), % of glial in mitosis (PH3-positive Glia /Total glial cells*100) (E), and % of glia cells incorporating EDU (F).

**Figure S9.**
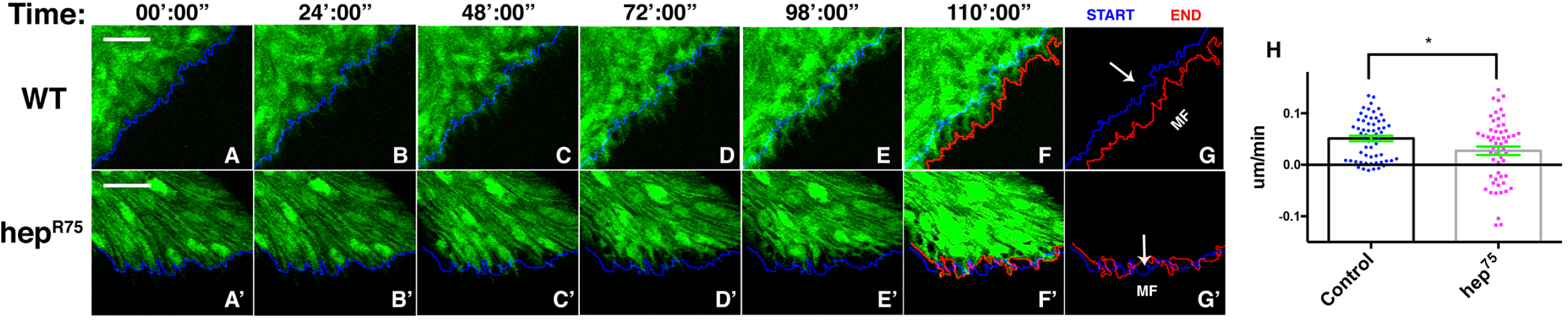
Glial motility is affected in *hep^R75^* mutants discs. Detailed frames from *in vivo* time-lapse analysis (video 3 and 4) of eye discs. Glial cells are labelled with *UAS-GFP* using *repo*-*Gal4*. (A-G) The migration of glial cells in control eye discs is coordinated and unidirectional, with a net displacement towards the location of the morfogenetic furrow (MF). (A’-G’) In *hep^r75^* mutant eye discs glial motility is strongly altered, therefore, most of glial cells remain in the same position, or they slightly advance. (H) The graph shows glial speed as defined as the distance covered (micrometres) by individual glial cells during the time-lapse analysis (minutes) with respect to the location of the MF. The blue lines indicate the start position of glial cells at the beginning of the time-lapse analysis, whereas red line indicates the most anterior position of glial cells at the end of the analysis. Arrows indicate the direction of the MF. Scale Bar 20 um

**Figure S10.**
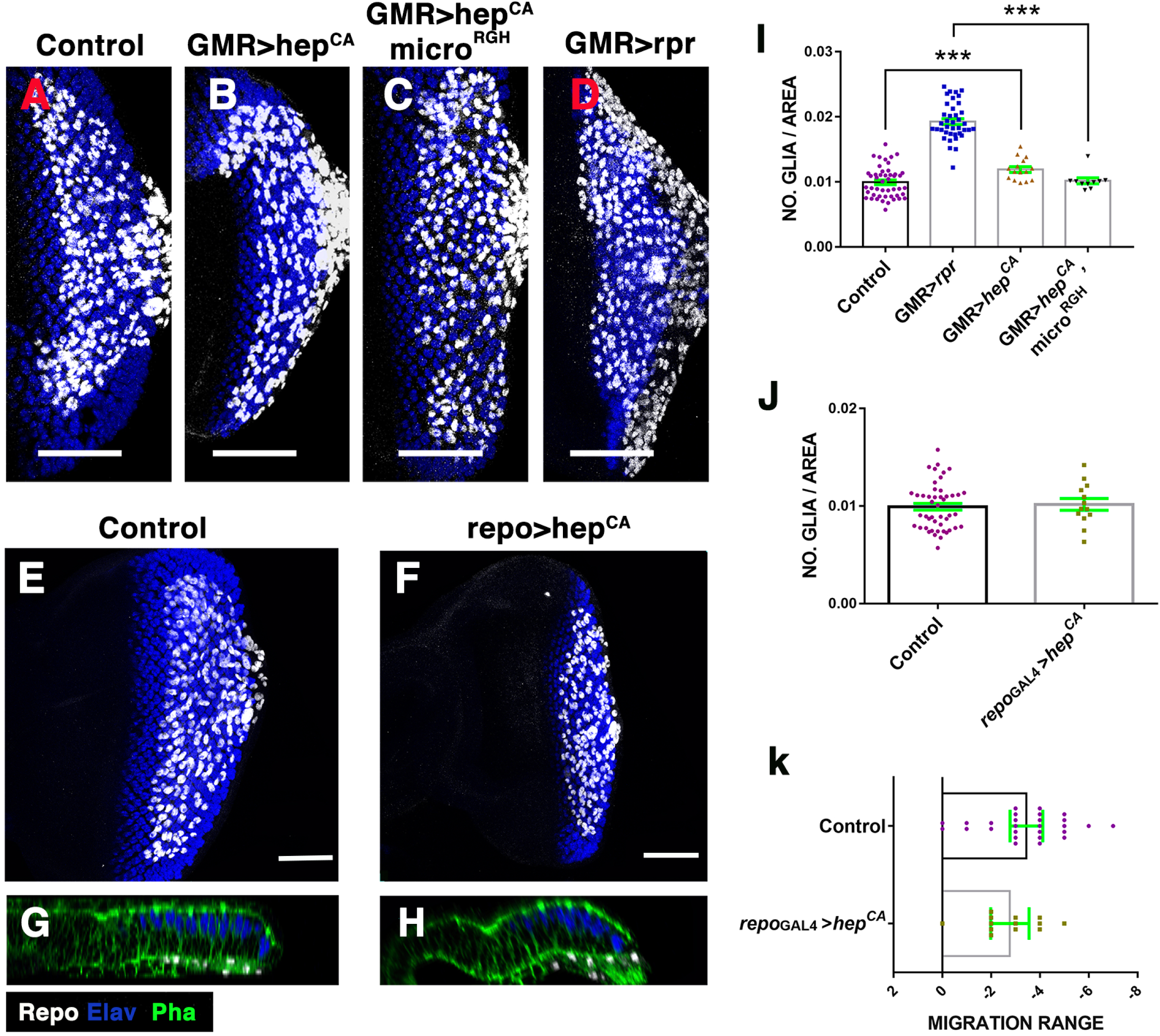
Overexpression of *hep^CA^* does not increase the number of glial cells in the eye discs. (A-F) Eye imaginal discs stained with anti-Repo (white) and anti-Elav (blue). Control (*GMR*-*Gal4 tub*-*Gal80^TS^*/*UAS*-GFP) (A), *GMR-Gal4 tub-Gal80^TS^/+; UAS-hep^CA^/+* (B), *GMR-Gal4 tub-Gal80^TS^/+; UAS-hep^CA^/micro^RGH^* (C) and *UAS-rpr/+GMR-Gal4 tub-Gal80^TS^/+ (*D), Control *tub-Gal80^TS^; repo-Gal4* (E), and *tub-Gal80^TS^/+; UAS-hep^CA^/repo-Gal4* (F). (G-H) X–Z projection show a cross-section of the eye discs epithelium perpendicular to the furrow of the discs shown in E-F. (I-J) Graphs show glial cell density (n° glial cells/ area) of the discs shown in A-D (I), and of the discs shown in E-F (J). (K) Graph shows the relative position of the anterior border of glial migration with respect to the anteriormost row of photoreceptors (0 indicates the position of this row) in control *tub-Gal80^TS^; repo-Gal4* and *tub-Gal80^TS^/+; UAS-hep^CA^/repo-Gal4* eye discs.

**Figure S11.**
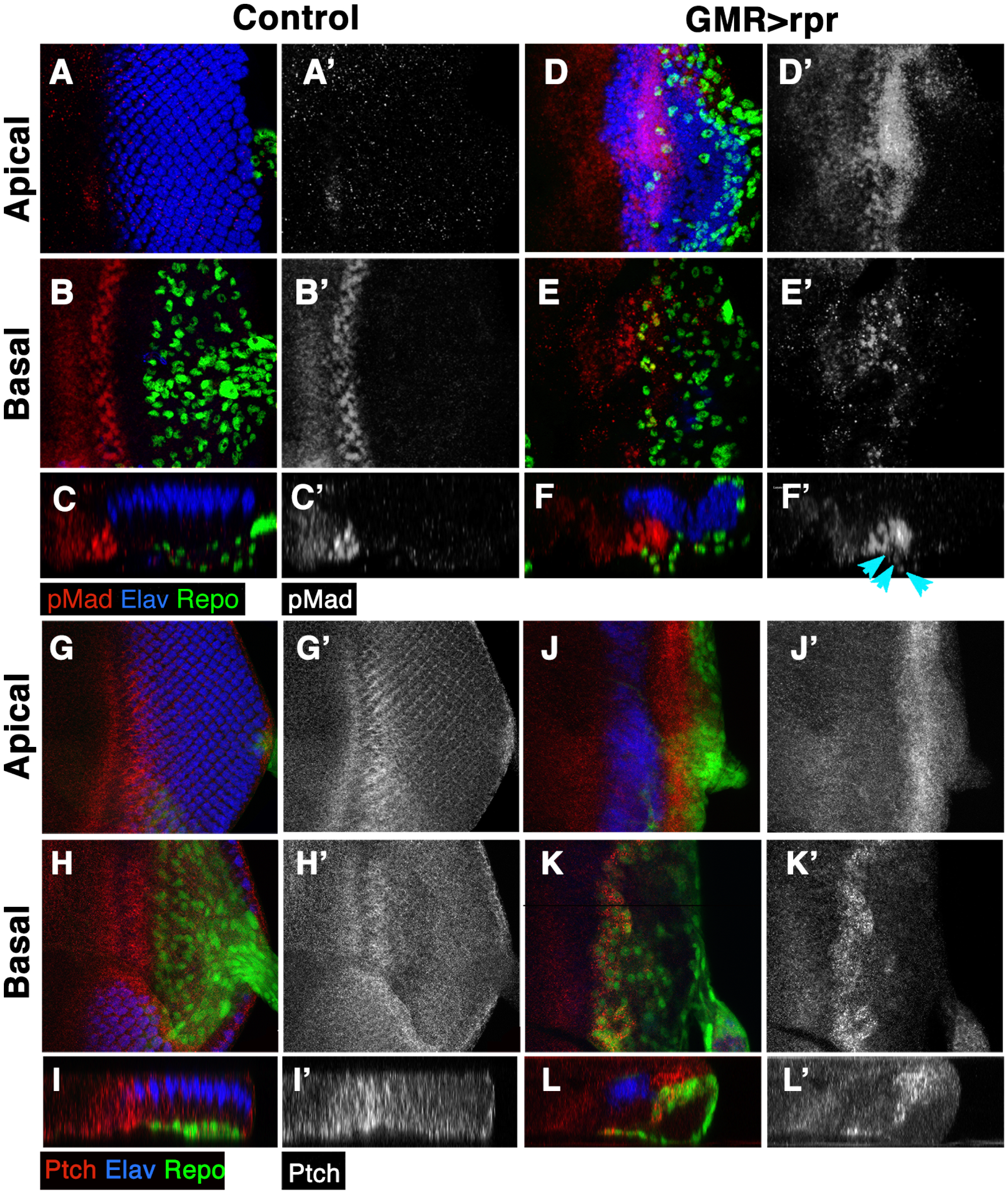
Expression of pMad and Patch increase in damaged discs. **(A-D)** Third instar eye discs stained with anti-Repo (green), anti-Elav (blue) and anti-pMad (in red A-B and D-F and in white A’-C’ and D’-F’) to reveal the activity of *dpp* signalling reporter (A-F’) or anti-Patch (G-L’). Control (*GMR-Gal4 tub-Gal80^TS^/+*) (A-C and G-I), and damaged disc *UAS*-*rpr*; *GMR*-*Gal4 tub*-*Gal80^TS^*/+ (D-F and –J-L’). Apical/middle layers (A-A’, D-D’, G-G’ and J-J’) and Basal layers (B-B’, E-E’, H-H’ and K-K’) of the eye disc epithelium. Cross-sections (C-C’, F-F’, I-I’ and L-L’) perpendicular to the furrow of the eye discs shown above. (A-F’) In control discs pMad is expressed in a band of cells along the furrow. However, in injured discs the expression of this factor increased in most of the cells behind the MF (D--F’), compared D-F’ to control discs A-C’. We also observed that the expression of pMad increases in some glial cells (blue arrowheads in F’). The Ptc expression is located at high levels in a band along the MF of the control discs (G-I’). In damaged discs Ptc signal was strong in most of the retinal and glial cells (J-L’).

**Figure S12.**
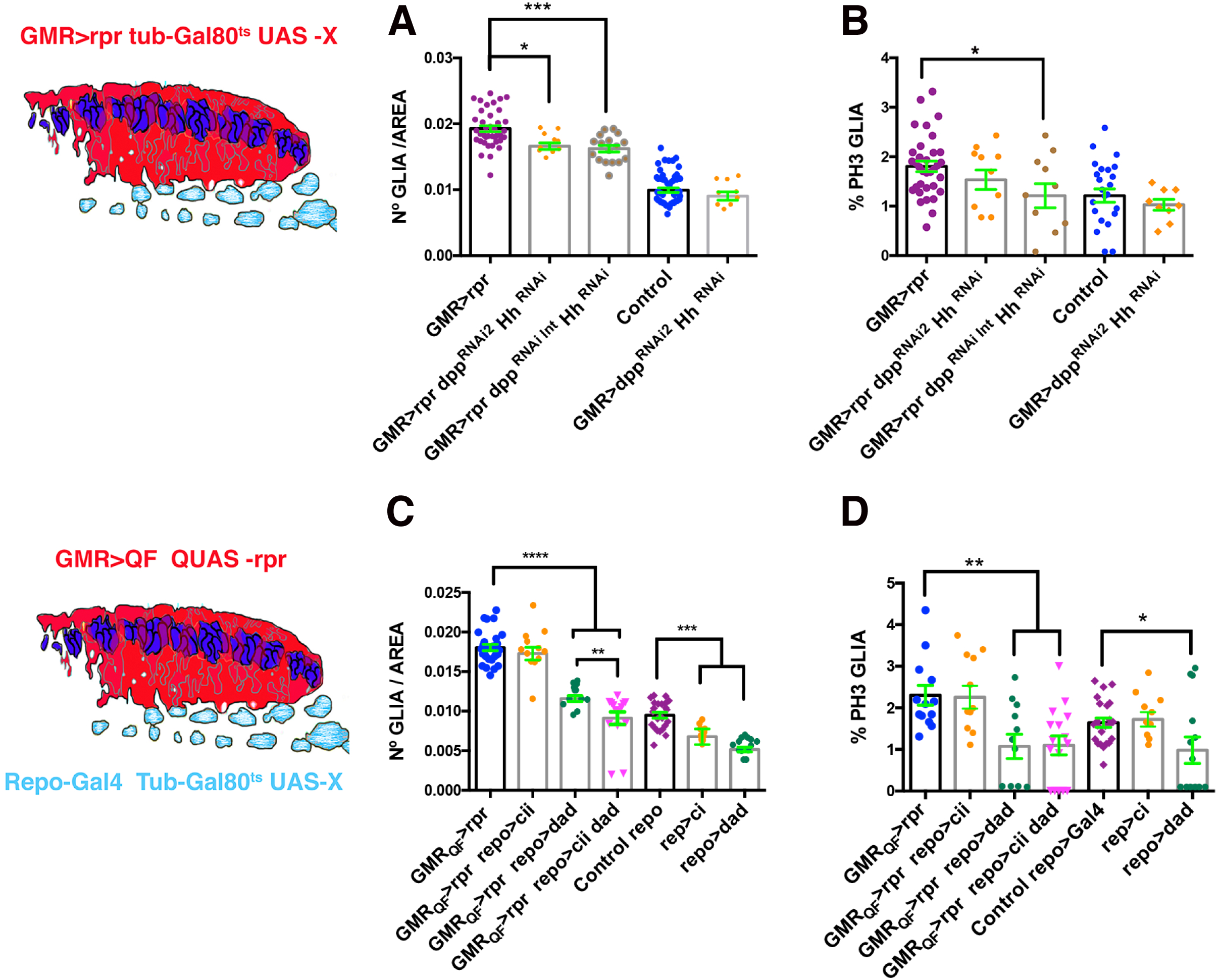
The down-regulation of *dpp* and *hh* signalling reduce glial response. The Schematic illustration on the left represents a transverse section of an eye disc where the region marked in red (*expression domain of GMR-Gal4)* corresponds to the area of the discs that has been damaged at the same time that *dpp* and/or *hh* signalling were depleted. (A-B) Graphs show glial cell density (A) and % of mitotic glia cells of: (*UAS-rpr; GMR-Gal4 tub-Gal80^ts^*) (*UAS-rpr/+; GMR-Gal4 tub-Gal80^ts^/ UAS-dpp^RNAi2^; UAS-hh^RNAi^*) (*UAS-rpr/+; GMR-Gal4 tub-Gal80^ts^/ UAS-dpp^RNAiint^; UAS-hh^RNAi^*) damaged discs, control undamaged *(GMR-Gal4 tub-Gal80^ts^*/+) (*GMR-Gal4 tub-Gal80^ts^/ UAS-dpp^RNAi2^*; *UAS-hh^RNAi^*). (C-D) Effects of the down-regulation of *dpp* and *hh* signalling in glial cells in damaged discs. In the schematic illustration on the left is indicated in red the region that has been damaged (*GMR-QF*; *QUAS-rpr*) and in light blue glial cells. (C-D) Graphs show glial cell density (C) and % of mitotic glia cells (D) of discs : *(GMR-QF; tub-Gal80^ts^*; *repo-Gal4 QUAS-rpr) (GMR-QF; tub-Gal80^ts^/+ repo-Gal4 QUAS-rpr/ UAS-ci^RNAi^)* (*GMR-QF; tub-Gal80^ts^/UAS-dad*; *repo-Gal4 QUAS-rpr/+)* (*GMR-QF; tub-Gal80^ts^/UAS-dad*; *repo-Gal4 QUAS-rpr/UAS-ci^RNAi^)* (*tub-Gal80^ts^; repo-Gal4, control)* (*tub-Gal80^ts^*; *repo-Gal4 /UAS-ci^RNAi^*) (*tub-Gal80^ts^*; *repo-Gal4/UAS-dad)*

**Figure S13.**
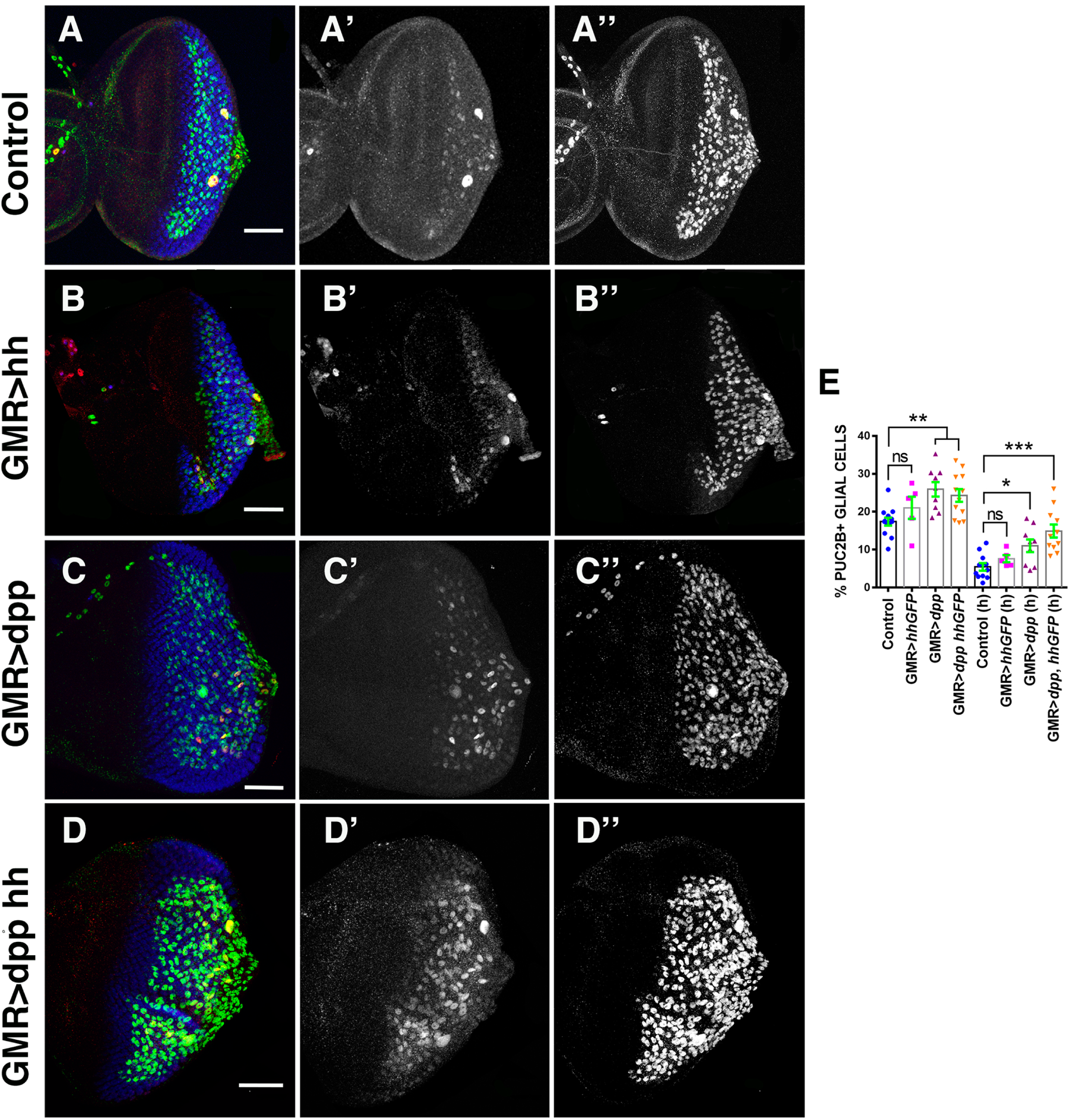
Over-expression of *dpp* in the retina region induces activation of JNK signalling in glial cells. (A-D’’) Third instar eye discs stained with anti-Elav (blue), anti-Repo (green in A-D and white A’’-D’’) and anti-B-galactosidase (red in A-D and white A’-D’) to reveal the activity of the *puc2b-LacZ* reporter, in control *(GMR-Gal4 tub-Gal80^ts^/+; puc2b-LacZ/+*) (A-A’’), discs over-expressing *UAS-hh* (*GMR-Gal4 tub-Gal80^ts^/UAS-hh; puc2b-LacZ*, B-B’’), discs over-expressing *UAS-dpp* (*GMR-Gal4 tub-Gal80^ts^/+; puc2b-LacZ/UAS-dpp*, C-C’’), and discs over-expressing simultaneously *UAS-dpp and UAS-hh* (*GMR-Gal4 tub-Gal80^ts^/UAS-hh; puc2b-LacZ/UAS-dpp*, D-D’’). (E) Graph shows % of glial cells expressing *puc2b-LacZ* at normal) and at high levels (h).

**Figure S14.**
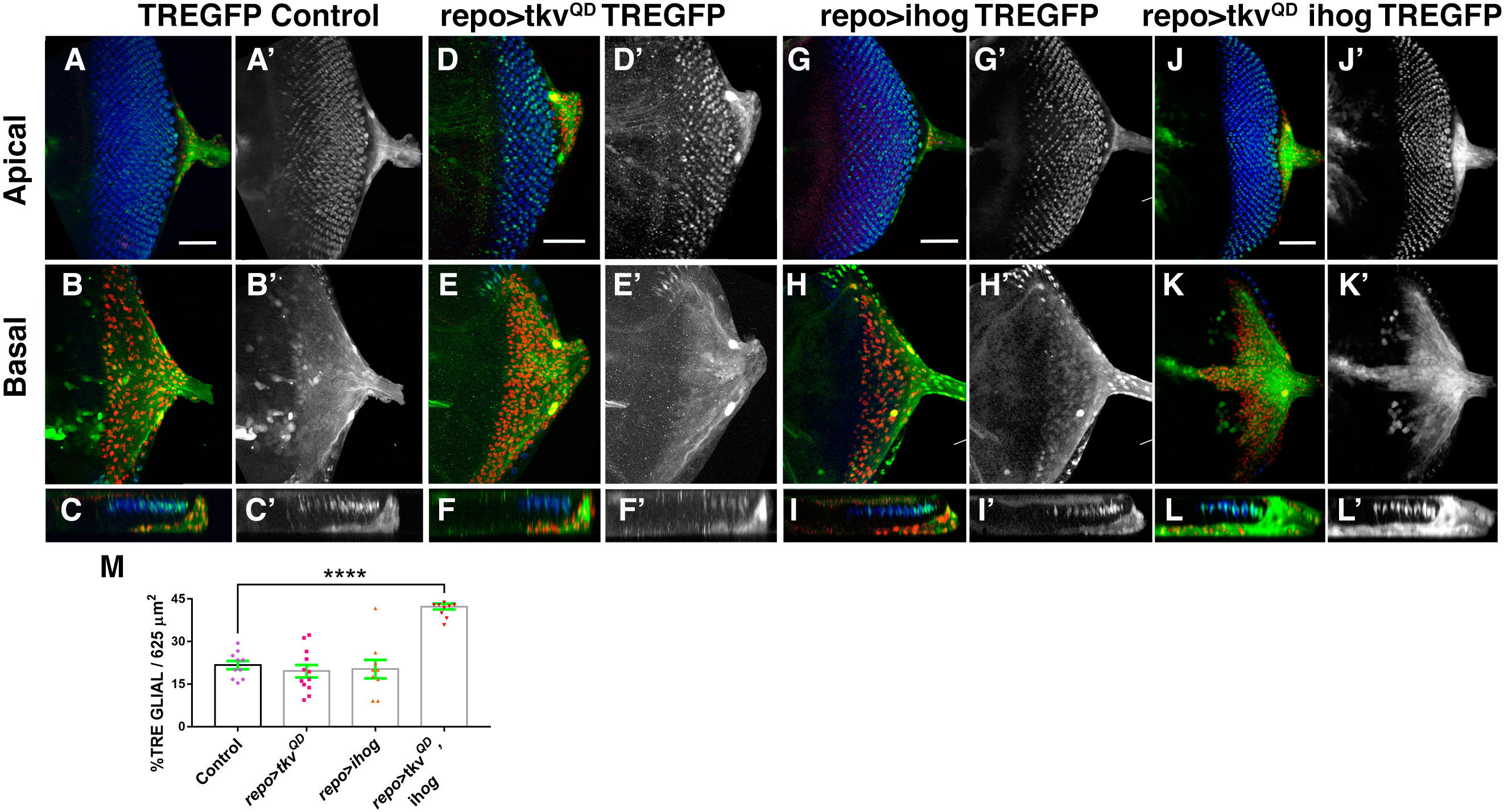
The activation of *dpp* and *hh* signalling in glial cells induce *TRE-GFP* over-expression in glial cells. (A-I’) Third instar eye discs stained with anti-Repo (red) and anti-Elav (blue). The *TRE-GFP* expression (green A-B, D-E, G-H and J-K and grey in A’-B’, D’-E’, G’-H’, and J’-K’) in control *(tub-Gal80^TS^/TRE-GFP; repo-Gal4*) (A-C’), *tub-Gal80^TS^/TRE-GFP; repo-Gal4/ UAS-tkv^QD^* (D-F’), *tub-Gal80^TS^/TRE-GFP; repo-Gal4 UAS-ihog/+* (G-I’) and *tub-Gal80^TS^/TRE-GFP; repo-Gal4 UAS-ihog/UAS-tkv^QD^,* eye discs (J-L’). Apical (A-A’, D-D’, G-G’ and J-J’), and Basal (B-B’, E-E’, H-H’, K-K’) layers. The X–Z projections below each panel show a cross-section of the epithelium perpendicular to the furrow. Bar charts show the average % of glial cells expressing *TRE-GFP*.

**Video 1**

Confocal timelapse imaging of damaged *GMR-QF; UAS-GFP Mz97-Gal4; QUAS-rpr* eye disc*. Mz97-Gal4* driving expression of *UAS-GFP* reveals wrapping glial cells WG extend large processes towards the damaged region. The timelapse covers a period of 160 minutes. Single frames for this video are shown in Figure 10J

**Video 2**

Confocal timelapse imaging of control *UAS-GFP Mz97-Gal4* eye disc*s*. *Mz97-Gal4* driving expression of *UAS-GFP* reveals wrapping glial cells. The timelapse covers a period of 160 minutes.

**Video 3**

Time lapse imaging showing the motility of glial cells in control eye discs. Glial cells are labelled with *UAS-GFP* using *repo*-*Gal4*. The timelapse covers a period of 140 minutes. Single frames for this video are shown in Figure S9

**Video 4**

Time lapse imaging showing the motility of glial cells in *hep^r75^*; *repo*-*Gal4 UAS-GFP* mutant eye discs. Most of glial cells remain in the same position, or they slightly advance. The timelapse covers a period of 140 minutes. Single frames for this video are shown in Figure S9B

**Video 5**

Confocal timelapse imaging of damaged *GMR-QF/UAS-bsk^DN^; UAS-GFP Mz97-Gal4; QUAS-rpr* eye disc*. Mz97-Gal4* drives *UAS-bsk^DN^* and *UAS-GFP* expression to reveal wrapping glial cells. To difference to WGs shown in video 2, WGs deprived of JNK signalling do not produce large cellular processes. The timelapse covers a period of 160 minutes. Single frames for this video are shown in Figure 10J

